# The RNA m^6^A reader YTHDF2 controls NK cell anti-tumor and anti-viral immunity

**DOI:** 10.1101/2021.04.26.441557

**Authors:** Shoubao Ma, Jiazhuo Yan, Tasha Barr, Jianying Zhang, Zhenhua Chen, Li-Shu Wang, Joseph C. Sun, Jianjun Chen, Michael A Caligiuri, Jianhua Yu

## Abstract

*N*^6^-methyladenosine (m^6^A) is the most prevalent post-transcriptional modification on RNA. NK cells are the predominant innate lymphoid cells that mediate anti-viral and anti-tumor immunity. However, whether and how m^6^A modifications affect NK cell immunity remains unknown. Here, we discover that YTHDF2, a well-known m^6^A reader, is upregulated in NK cells upon activation by cytokines, tumors, and cytomegalovirus infection. *Ythdf2* deficiency in NK cells impairs NK cell anti-tumor and anti-viral activity *in vivo*. YTHDF2 maintains NK cell homeostasis and terminal maturation, correlating with modulating NK cell trafficking and regulating Eomes, respectively. YTHDF2 promotes NK cell effector function and is required for IL-15-mediated NK cell survival and proliferation by forming a STAT5-YTHDF2 positive feedback loop. Transcriptome-wide screening identifies *Tardbp* to be involved in cell proliferation or survival as a YTHDF2-binding target in NK cells. Collectively, we elucidate the biological roles of m^6^A modifications in NK cells and highlight a new direction to harness NK cell anti-tumor immunity.

## Introduction

Natural killer (NK) cells are the predominant innate lymphoid cells (ILC) that mediate anti-viral and anti-tumor immunity (Spits et al., 2016). They recognize virus-infected and cancerous cells through their multiple surface-expressed activating and inhibitory receptors and kill them via a cytotoxic effect (Sun and Lanier, 2011). They can also produce a distinct set of cytokines such as IFN-γ, TNF-α, IL-10, or chemokines such as MIP-1α and -β and RANTES, which can further elicit adaptive immune responses (Spits et al., 2016). Together, the multifunctional activities of NK cells help to eliminate susceptible targets and amplify inflammatory responses against viruses and cancer.

As the most prevalent post-transcriptional modification on mammalian mRNA, the *N*^6^-methyladenosine (m^6^A) modification is emerging as a widespread regulatory mechanism that controls gene expression in diverse physiological processes (Yue et al., 2015). However, how m^6^A methylations regulate innate and adaptive cell-mediated immunity remains to be fully understood and, up until this report, has remained unknown in NK cells. Recently, chimeric antigen receptors (CARs) have been shown to re-direct NK cells toward tumor cells expressing a corresponding antigen, creating opportunities to fight against cancer (Chen et al., 2016; Chu et al., 2014; Han et al., 2015; Liu et al., 2020; Tang et al., 2018; Yilmaz et al., 2020). Therefore, clearly defining the role of m^6^A modification in NK cells will not only greatly improve our understanding of RNA modifications as a novel and critical layer of post-transcriptional gene regulation that controls innate immune cell functions but may also provide us a new opportunity to enhance NK cell effector function and survival for cancer immunotherapy.

The m^6^A methyltransferases (“writers”, i.e., METTL3 and METTL14) and demethylases (“erasers”, i.e., FTO and ALKBH5) dynamically control the m^6^A methylation landscape within the nucleus (Shi et al., 2019). The m^6^A reader proteins (YTH domain-containing family (YTHDF) proteins, YTHDF1, YTHDF2, and YTHDF3; and insulin-like growth factor 2 mRNA-binding (IGF2BP) proteins, IGF2BP1, IGF2BP2, and IGF2BP3) preferentially bind to the methylated RNA and mediate specific functions, including promoting the translation or affecting stability of m^6^A-modified mRNAs (Huang et al., 2018; Shi et al., 2019; Wang et al., 2014; Wang et al., 2015). Recent studies have shown that m^6^A methylation is involved in adaptive and innate immune cell-mediated immunity (Shulman and Stern-Ginossar, 2020).

Deletion of m^6^A “writer” protein METTL3 in mouse T cells disrupts cell homeostasis and differentiation by targeting the IL-7/SOCS/STAT5 pathway (Li et al., 2017). METTL3 maintains Treg cell suppressive functions through IL-2-STAT5 signaling (Tong et al., 2018). RNA m^6^A methylation plays an essential role in early B cell development (Zheng et al., 2020). A recent report shows that METTL3-mediated mRNA m^6^A methylation promotes dendritic cell (DC) activation (Wang et al., 2019a). m^6^A-modified mRNAs encoding lysosomal cathepsins can be recognized by YTHDF1 in DCs, thereby suppressing the cross-priming ability of DCs and inhibiting anti-tumor immune responses (Han et al., 2019). m^6^A modifications also control the innate immune response to virus infection (Liu et al., 2019; Rubio et al., 2018; Winkler et al., 2019). However, whether and how m^6^A modifications affect the NK cell-mediated immune response to tumor cells and virus has not been reported.

YTHDF2 is a well-recognized m^6^A reader that acts by specifically recognizing and binding to m^6^A-containing RNAs and promoting degradation of target transcripts (Wang et al., 2014). According to the database of BioGPS (Wu et al., 2013) and our preliminary data, murine NK cells express YTHDF2 at a high level, while its role in regulating NK cells is unknown. This motivates us to study YTHDF2 in NK cells using a conditional knockout approach. We show that depletion of *Ythdf2* in mouse NK cells significantly impaired NK cell anti-tumor and antiviral immunity. Moreover, YTHDF2 controlled NK cell homeostasis, maturation, and survival at a steady state. Thus, YTHDF2 or m^6^A in general plays multifaceted roles in regulating NK cells.

## Results

### YTHDF2 is upregulated in murine NK cells by IL-15, MCMV infection, and tumor progression

To study the role of m^6^A modifications in NK cells, we first screened the expression levels of m^6^A “writers”, “erasers”, and “readers” in murine NK cells by using BioGPS database (http://biogps.org). Accordingly, the expression of *Ythdf2* mRNA was the highest among the other m^6^A enzymes and readers (Fig. 1 A). Interestingly, we found that NK cells also have constitutive mRNA and protein expression of *Ythdf2* at high levels compared to most other immune cells, including B cells, macrophages, and DCs (Fig. 1 B and Fig. S1 A). NK cells can be activated by IL-15, which is a key regulator of NK cell homeostasis and survival (Becknell and Caligiuri, 2005). Leveraging the GEO database, we found that IL-15-activated NK cells have higher mRNA levels of *Ythdf2* compared to other m^6^A enzymes and readers that we tested (Fig. 1 C, analyzed from GSE106138). Consistent with our analysis of the BioGPS data, our real-time quantitative PCR (qPCR) and immunoblotting showed that IL-15 activation of NK cells significantly upregulated *Ythdf2* at both the mRNA and protein levels, and that *Ythdf2* seemed to have the highest expression levels in IL-15-activated NK cells compared to other m^6^A enzymes and readers that we tested (Fig. 1, D-F; and Fig. S1 B). The protein levels of YTHDF2 were also significantly upregulated in NK cells from IL-15Tg mice that we previously generated compared to wild-type control (Fehniger et al., 2001) (Fig. S1 C).

**Figure 1.**
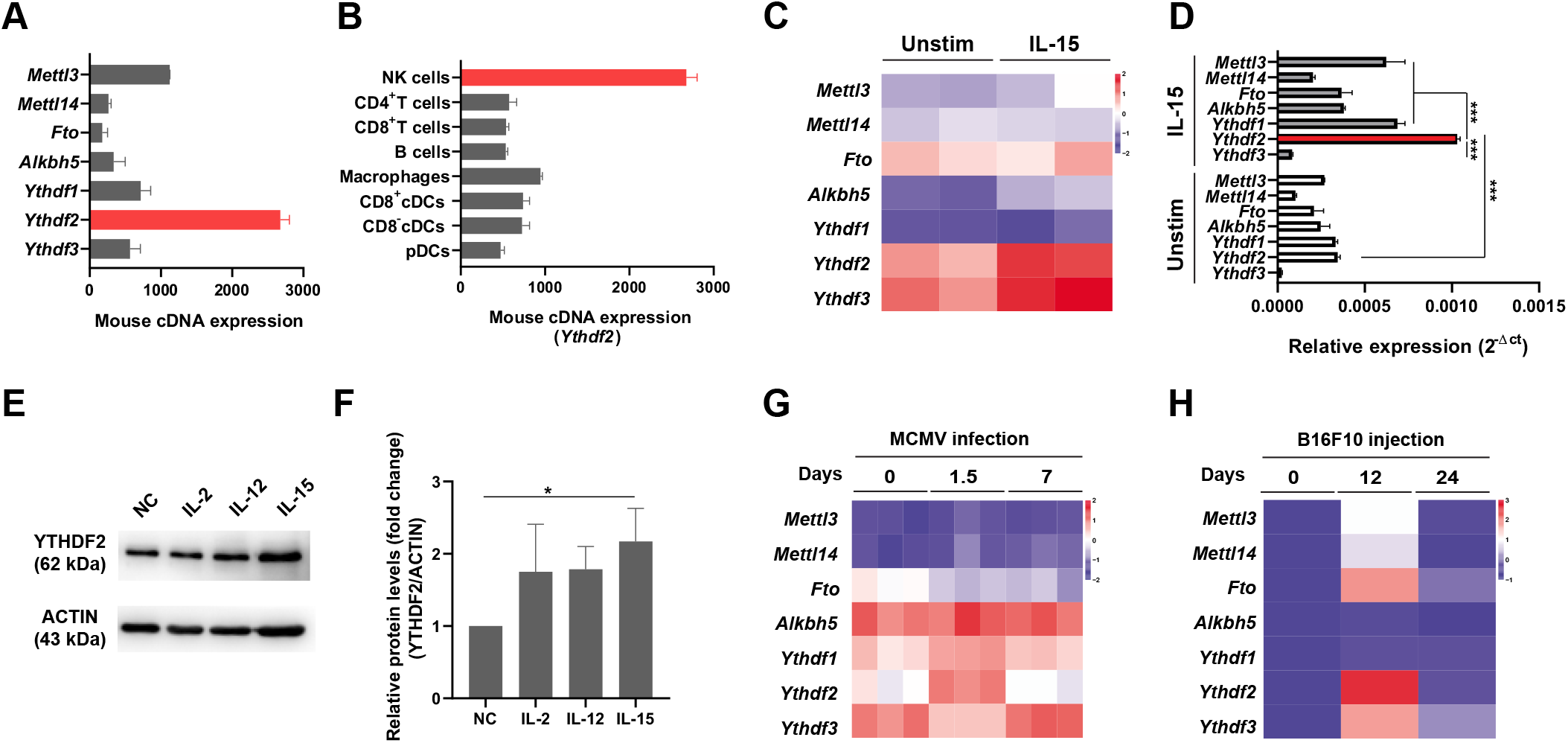
YTHDF2 expression in murine NK cells in response to IL-15 stimulation, MCMV infection, and tumor progression. **(A-B)** The mRNA expression of m^6^A “writers” (*Mettl3, Mettl14*), “erasers” (*Fto, Alkbh5*), and “readers” (*Ythdf1, Ythdf2, Ythdf3*) in murine NK cells (**A**) and the mRNA levels of *Ythdf2* among immune cell subsets (**B)** were analyzed using the BioGPS online tool. **(C)** RNA-seq analysis of the online database GSE106138 showing the expression of m^6^A enzymes and readers in splenic NK cells cultured either in the presence or absence of IL-15 (10 ng/ml) overnight. **(D)** qPCR showing the expression of m^6^A enzymes and readers in splenic NK cells cultured either in the presence or absence of IL-15 (10 ng/ml) overnight. *18s rRNA* was used as a housekeeping gene for data normalization. Data are shown as mean ± SD (\*\*\**P* < 0.001; unpaired two-tailed *t*-test,). Average values from 3 replicates were calculated for each sample. The experiment was repeated three times independently. **(E-F)** Representative immunoblot (**E**) and quantified protein levels of YTHDF2 normalized to ACTIN (**F**) in splenic NK cells cultured in the presence or absence of IL-2 (100 U/ml), IL-12 (10 ng/ml), or IL-15 (10 ng/ml) overnight. Untreated NK cells were used as negative control (NC group). **(G)** RNA-seq analysis of online database GSE113214 showing the expression of m^6^A enzymes and readers in splenic NK cells during the MCMV infection. **(H)** NK cells were isolated from lung tissues of mice injected with B16-F10 at indicated time points. The expression levels of m^6^A enzymes and readers were examined by qPCR. Heatmap showing the expression m^6^A enzymes and readers in the isolated NK cells after B16F10 injection. *18s rRNA* was used as a housekeeping gene for data normalization. The expression of each gene is shown as a fold change from the data collected on day 0. Data are shown as mean ± SD and were analyzed by one-way ANOVA with Sidak posttest (**F**, \**P* < 0.05). Data are representative of at least two independent experiments.

NK cells are critical mediators of host immunity against viral infection and malignancies (Spits et al., 2016). We therefore evaluated the expression pattern of YTHDF2 in NK cells during murine cytomegalovirus (MCMV) infection. Using the GEO database, we found that *Ythdf2* was upregulated at day 1.5 post-infection in two independent databases (Fig. 1 G and Fig. S1 D). An immunoblotting assay confirmed that the protein levels of YTHDF2 were also upregulated at day 1.5 post-infection with MCMV (Fig. S1 E), indicating that YTHDF2 may play a critical role in NK cell-mediated anti-viral immunity. In addition to controlling virus infection, NK cells also contribute to anti-tumor immunosurveillance. We then examined the *Ythdf2* levels during tumor development. Using the B16F10 melanoma metastasis model, we found a reduction of NK cells in the lung at the late stage of tumor development (Fig. S1 F), which is consistent with a prior report (Cong et al., 2018). We found that the mRNA and protein levels of *Ythdf2* were significantly upregulated in NK cells at the early stage of tumor development (Fig. 1 H and Fig. S1 G). Taken together, these data demonstrate that *Ythdf2* is highly expressed in NK cells and is upregulated during viral infection and tumorigenesis, leading us to hypothesize that YTHDF2 plays a role in regulating NK cell defense against tumorigenesis and viral infection.

### YTHDF2 deficiency impairs NK cell anti-tumor immunity

To define the role of YTHDF2 in NK cell-mediated anti-tumor immunity, we used the CRISPR-Cas9 technology to generate *Ythdf2* floxed mice (Fig. S1 H). We then generated NK cell-specific conditional knockout mice (hereafter referred to as *Ythdf2*^ΔNK^ mice) by crossing *Ythdf2*^fl/fl^ mice with *Ncr1*-iCre mice (Narni-Mancinelli et al., 2011). Deletion of *Ythdf2* in NK cells was verified by qPCR and immunoblotting (Fig. S1, I and J). We then established a metastatic melanoma model by intravenous injection of B16F10 cells into *Ythdf2*^WT^ and *Ythdf2*^ΔNK^ mice. As shown in Fig. 2 A, Ythdf2^ΔNK^ mice displayed a much greater burden of metastatic nodules than that of *Ythdf2*^WT^ mice. We found a significant reduction in the percentage and absolute number of infiltrating NK cells in tumor tissues of *Ythdf2*^ΔNK^ mice compared to those observed in *Ythdf2*^WT^ mice (Fig. 2, B and C). Meanwhile, infiltrating NK cells from *Ythdf2*^ΔNK^ mice showed a significant decrease in the expression of IFN-γ, granzyme B, and perforin compared with those from *Ythdf2*^WT^ mice (Fig. 2, D-F). However, the percentages of CD4^+^T cells and CD8^+^T cells and their expression of IFN-γ were comparable between *Ythdf2*^WT^ mice and *Ythdf2*^ΔNK^ mice (Fig. S1, K-N), suggesting that YTHDF2 in NK cells is essential for controlling tumor metastases. To confirm the cell-intrinsic requirement of YTHDF2 for NK cell-mediated anti-tumor immunity, we adoptively transferred an equal number of NK cells from *Ythdf2*^ΔNK^ mice or *Ythdf2*^WT^ mice into *Rag2*^−/−^*Il2rg*^−/−^ mice which lack T, B, and NK cells, one day prior to an injection of B16F10 tumor cells (Fig. 2 G). We found a significantly increased incidence of tumor metastases in mice transferred with *Ythdf2*^ΔNK^ NK cells compared with mice injected with *Ythdf2*^WT^ NK cells (Fig. 2 G). Similarly, in this model, we also found a significant reduction in the percentage and the absolute number of infiltrating *Ythdf2*^ΔNK^ NK cells (Fig. 2, H and I), as well as a decrease in the expression of IFN-γ, granzyme B, and perforin in mice that received adoptively transferred *Ythdf2*^ΔNK^ NK cells compared to those that received *Ythdf2*^WT^ NK cells (Fig. 2, J-L). These data indicate a cell-intrinsic role of YTHDF2 in the regulation of NK cell anti-tumor immunity.

**Figure 2.**
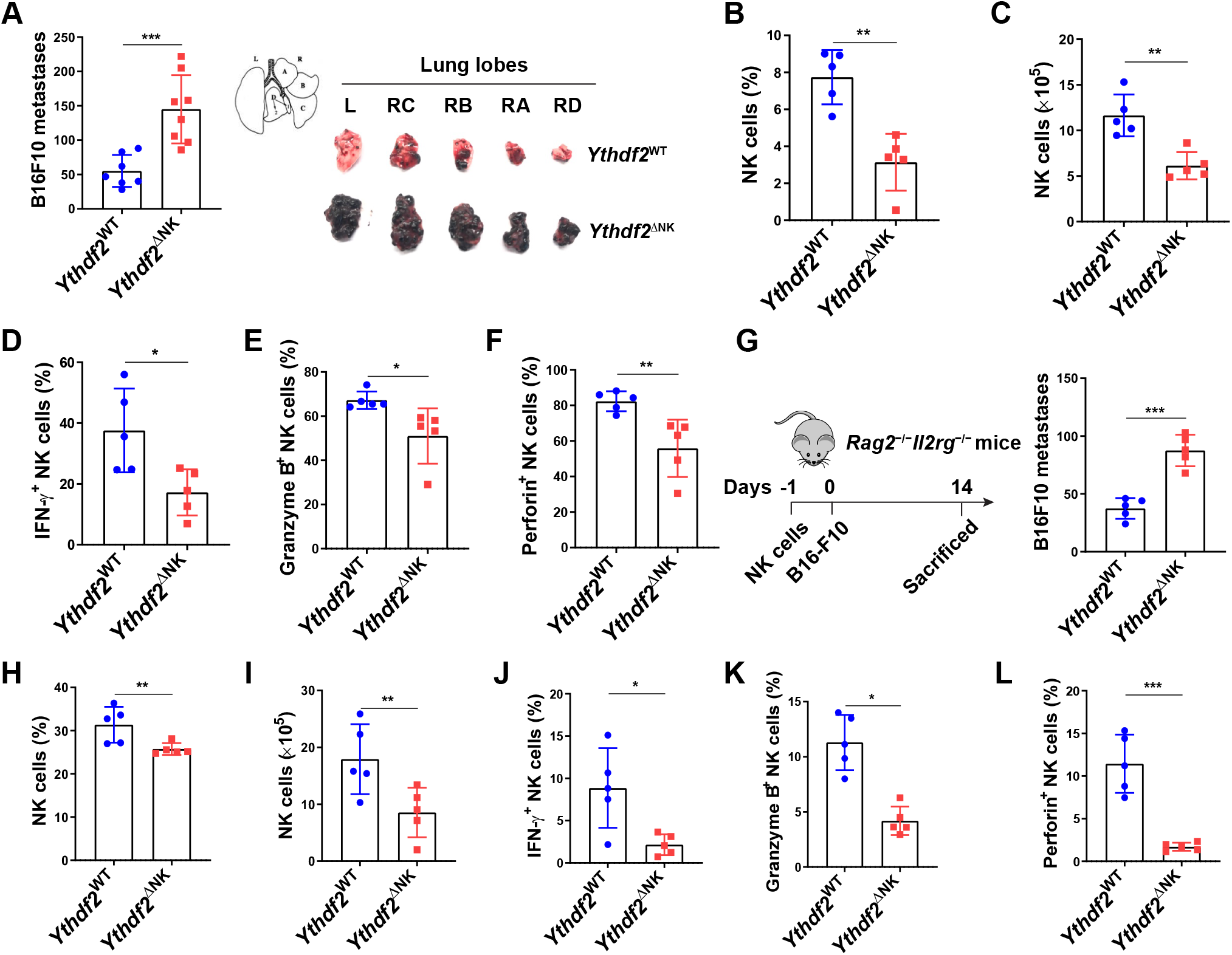
Requirement for YTHDF2 in NK cell anti-tumor immunity. **(A-F)** *Ythdf2*^WT^ and *Ythdf2*^ΔNK^ mice were i.v. injected with B16F10 cells (1 × 10^5^). Fourteen days after injection, the mice were euthanized for post-mortem analysis. Quantification of total metastatic nodules in the lung from *Ythdf2*^WT^ (n = 7) and *Ythdf2*^ΔNK^ mice (n = 8); gross morphology of individual lung lobes is shown on the right **(A)**. The percentage and the absolute number of infiltrating NK cells in lung tissues from *Ythdf2*^WT^ and *Ythdf2*^ΔNK^ mice (n = 5 mice per group, at least two independent experiments) **(B-C)**. IFN-γ, granzyme B, and perforin expression in the lung-infiltrated NK cells from *Ythdf2*^WT^ and *Ythdf2*^ΔNK^ mice (n = 5 mice per group, at least two independent experiments) **(D-F)**. **(G-L**) 1 × 10^6^ IL-2 expanded NK cells from *Ythdf2*^WT^ and *Ythdf2*^ΔNK^ mice were i.v. injected into *Rag2*^−/−^*Il2rg*^−/−^ mice. One day later, B16F10 cells (1 × 10^5^) were i.v. injected into mice. Fourteen days after injection, mice were euthanized for post-mortem analysis. Quantification of total metastatic nodules in the lung from *Ythdf2*^WT^ and *Ythdf2*^ΔNK^ mice (n = 5 mice per group, at least two independent experiments) **(G)**. The percentage and the absolute number of infiltrating NK cells in the lung tissues (n = 5 mice per group, at least two independent experiments) **(H-I)**. IFN-γ, granzyme B, and perforin expression in the lung-infiltrated NK cells from *Ythdf2*^WT^ and *Ythdf2*^ΔNK^ mice (n = 5 mice per group, at least two independent experiments) **(J-L)**. Each symbol represents an individual mouse. Data are shown as mean ± SD and were analyzed by unpaired two-tailed *t*-test (**A-L**, \**P* < 0.05, \*\**P* < 0.01, \*\*\**P* < 0.001).

### YTHDF2 is required for the anti-viral function of NK Cells

To determine whether *Ythdf2* deficiency affects the anti-viral activity of NK cells, we injected 2.5×10^4^ plaque-forming units (pfu) of MCMV into *Ythdf2*^WT^ mice and *Ythdf2*^ΔNK^ mice. The results showed that *Ythdf2*^ΔNK^ mice were more susceptible to MCMV infection, as depicted by significant weight loss and increased viral titers in blood, spleen, and liver compared to *Ythdf2*^WT^ mice (Fig. 3, A and B; and Fig. S2, A and B). We also observed a significant reduction in the percentage and the absolute number of total NK cells in spleen and blood of *Ythdf2*^ΔNK^ mice compared to those in *Ythdf2*^WT^ mice post-infection (Fig. 3, C and D; and Fig. S2, C and D). Further analysis showed that NK cells from *Ythdf2*^ΔNK^ mice had significantly lower expression of Ki67 than that of *Ythdf2*^WT^ mice (Fig. 3, E-G). However, NK cell viability was similar in *Ythdf2*^WT^ mice and *Ythdf2*^ΔNK^ mice, as shown by Annexin V staining (Fig. S2 E). These data indicate the deficiency of YTHDF2 in NK cells results in a defect in cell proliferation rather than cell survival during viral infection. NK cells inhibit MCMV infection through the activating receptor Ly49H and Ly49D and are characterized by a perforin- or IFN-γ-mediated anti-viral response (Arase et al., 2002; Lee et al., 2009; Loh et al., 2005; Orr et al., 2010; Sumaria et al., 2009). We found that *Ythdf2*^ΔNK^ mice had significantly reduced Ly49H^+^ and Ly49D^+^ NK cells in spleen and blood compared to that of *Ythdf2*^WT^ mice post-infection (Fig. 3, H-J; and Fig. S2, F-K). Further analysis demonstrated that although per percentage, only Ly49D^+^Ly49H^+^ cells showed a difference in *Ythdf2*^ΔNK^ mice compared to that of *Ythdf2*^WT^ mice, the absolute cell numbers of Ly49D^-^Ly49H^+^ NK cells, Ly49D^+^Ly49H^-^ NK cells, and Ly49D^+^Ly49H^+^ NK cells in the spleen and blood were all significantly decreased in *Ythdf2*^ΔNK^ mice compared to that of *Ythdf2*^WT^ mice post-infection (Fig. S2, L-W). Our data suggest that controlling MCMV infection by YTHDF2 seems to be mainly mediated by Ly49D^+^Ly49H^+^ NK cells. The granzyme B and IFN-γ production by NK cells in *Ythdf2*^ΔNK^ mice was comparable to that of *Ythdf2*^WT^ mice (Fig. S2, X and Y). We found significantly reduced perforin production by *Ythdf2*^ΔNK^ mice compared to that of *Ythdf2*^WT^ mice in both spleen and blood 7 days post-infection (Fig. 3, K-M), indicating that YTHDF2 mainly affects perforin-mediated anti-viral activity against MCMV in NK cells. These data indicate that YTHDF2 is critical for NK cell expansion and effector function during MCMV infection.

**Figure 3.**
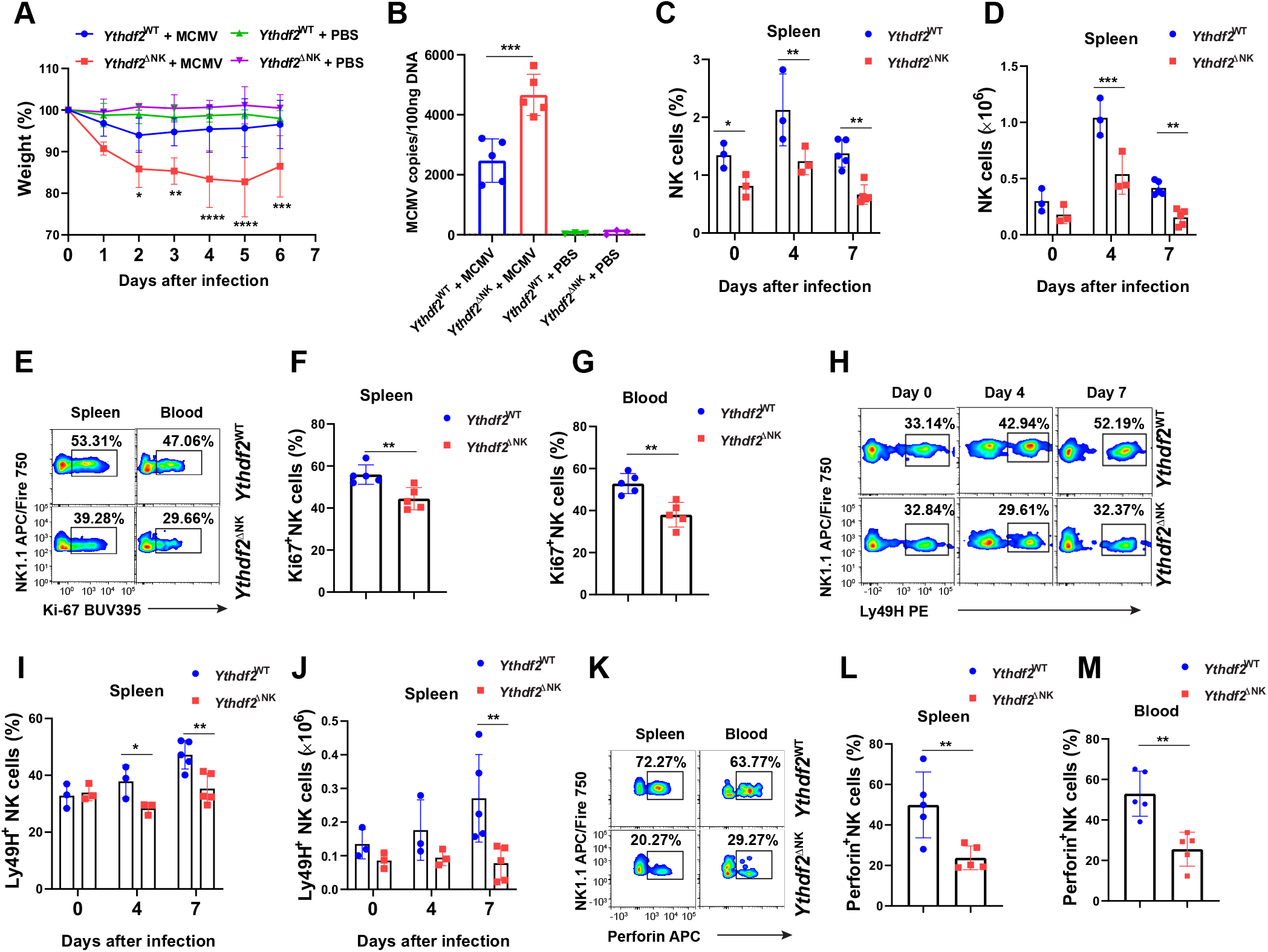
*Ythdf2*-deficient NK cells have impaired anti-viral functions. **(A)** 2.5 × 10^4^ plaque-forming units (pfu) of MCMV were injected i.p. into *Ythdf2*^WT^ and *Ythdf2*^ΔNK^ mice. Weight loss of *Ythdf2*^WT^ and *Ythdf2*^ΔNK^ mice at various times after infection is shown, presented as relative weights compared to pre-infection (n = 5 mice for MCMV groups, and n = 3 for PBS groups). **(B)** Viral titers in blood on day 7 post-infection were assessed by qPCR. Mice injected with PBS were used as control (n = 5 mice for MCMV groups; n = 3 for PBS groups). **(C-D)** The percentage and the absolute number of NK cells in the spleen from *Ythdf2*^WT^ and *Ythdf2*^ΔNK^ mice on days 0, 4, 7 post-infection with MCMV (n = 3 at days 0 and 4; n = 5 at day 7). **(E-G)** Representative plots (**E**) and quantification of Ki67 expression by NK cells in the spleen (**F**) and blood (**G**) from *Ythdf2*^WT^ and *Ythdf2*^ΔNK^ mice on day 7 post-infection (n = 5 mice per group). **(H-J)** Representative plots (**H**) and quantification of the percentage (**I**) and absolute number (**J**) of Ly49H^+^ NK cells in spleen from *Ythdf2*^WT^ and *Ythdf2*^ΔNK^ mice on days 0, 4, 7 post-infection with MCMV (n = 3 at days 0 and 4; n = 5 at day 7). **(K-M)** Representative plots (**K**) and quantification of perforin expression by NK cells in the spleen (**L**) and blood (**M**) from *Ythdf2*^WT^ and *Ythdf2*^ΔNK^ mice on day 7 post-infection (n = 5 mice per group). Data are shown as mean ± SD and were analyzed by unpaired two-tailed *t*-test (**F, G, L, M**), one-way ANOVA with Sidak post-test (**B**), or two-way ANOVA with Sidak post-test (**A, C, D, I, J**). \**P* < 0.05, \*\**P* < 0.01, \*\*\**P* < 0.001, \*\*\*\**P* < 0.0001. Data are representative of at least three independent experiments.

### YTHDF2 controls NK cell homeostasis and terminal maturation at a steady state

The above findings that *Ythdf2* deficiency in NK cells enhanced tumor metastases and impaired NK capacity to control MCMV infection encouraged us to investigate whether YTHDF2 was required for NK cell maintenance at a steady state. As shown in Fig. 4 A, the frequency and the absolute number of NK cells were significantly reduced in the peripheral blood, spleen, liver, and lung but not bone marrow (BM) in *Ythdf2*^ΔNK^ mice compared with *Ythdf2*^WT^ mice. However, there were no significant changes among common lymphoid progenitor (CLP), pre-NK cell progenitor (preNKP), and refined NKP (rNKP) in the BM (Fathman et al., 2011) between *Ythdf2*^WT^ and *Ythdf2*^ΔNK^ mice (Fig. S3 A), indicating that YTHDF2 may not affect NK cell early development in our model. To explore the potential mechanisms responsible for the decrease of NK cells in *Ythdf2*^ΔNK^ mice, we investigated cell proliferation, viability, and trafficking ability of NK cells after *Ythdf2* deletion at a steady state. The percentage of proliferating NK cells was comparable between *Ythdf2*^WT^ and *Ythdf2*^ΔNK^ mice, as evidenced by Ki67 staining (Fig. S3 B). The viability of NK cells was also equivalent between *Ythdf2*^WT^ and *Ythdf2*^ΔNK^ mice, as shown by Annexin V staining (Fig. S3 C). To check whether YTHDF2 affects egress of NK cells from BM to the periphery, *Ythdf2*^WT^ and *Ythdf2*^ΔNK^ mice were intravenously injected with an anti-CD45 antibody to mark immune cells, sacrificed after 2 min, and their BM cells were analyzed. This allowed us to quantify the number of NK cells in the sinusoidal versus parenchymal regions of the BM, an indicator of NK cell trafficking from BM to peripheral blood under a steady state (Leong et al., 2015). The results showed a significant reduction in the frequency of CD45^+^ NK cells in *Ythdf2*^ΔNK^ mice compared to *Ythdf2*^WT^ mice in the sinusoids (Fig. 4 B), indicating that *Ythdf2* deficiency impairs the egress of NK cells from bone marrow to the circulation system *in vivo*.

**Figure 4.**
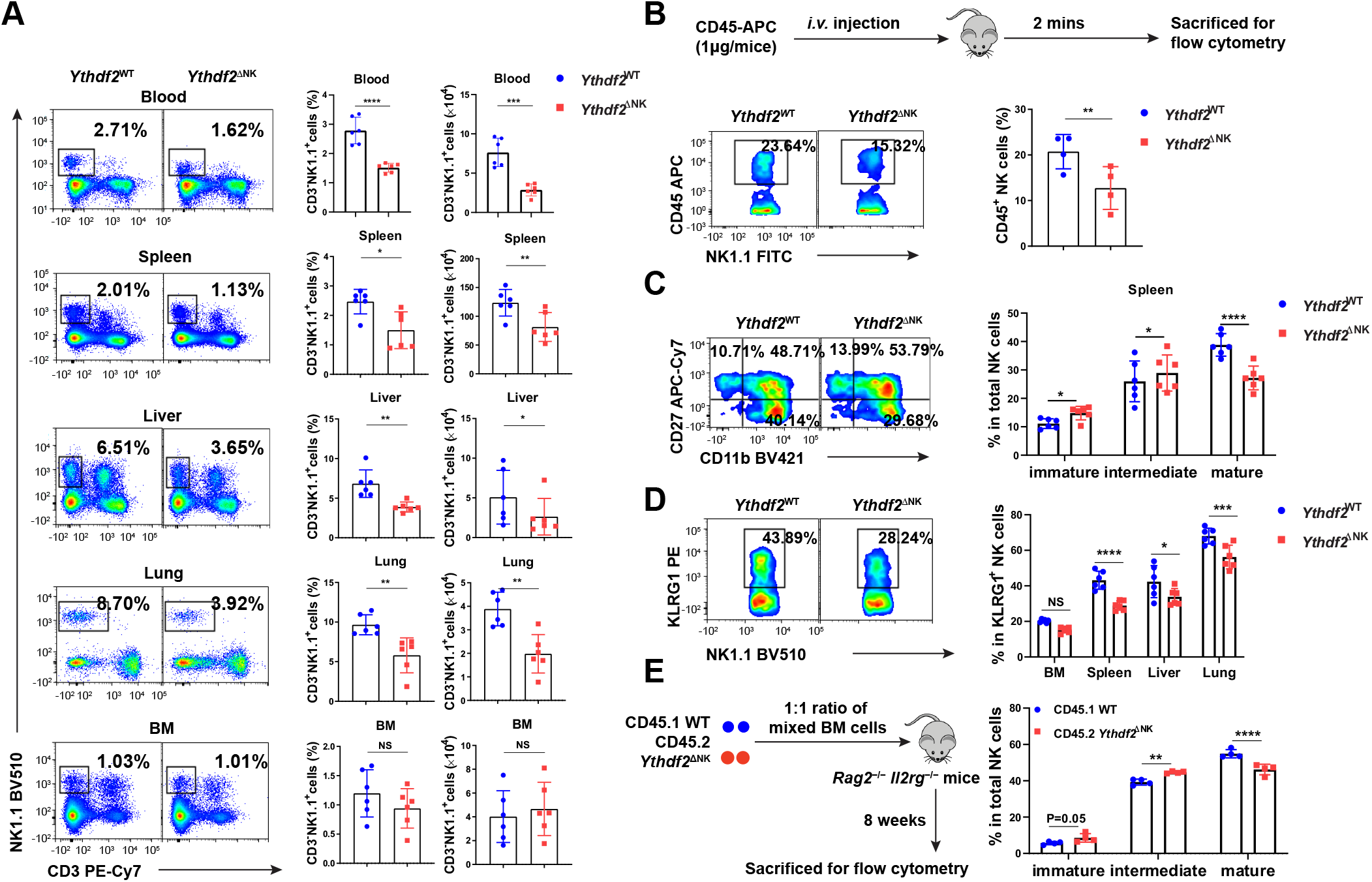
*Ythdf2* deficiency inhibits NK cell homeostasis and terminal maturation at a steady state. **(A)** Representative plots (left) and quantification (right) of the percentage and the absolute number of NK cells (CD3^-^NK1.1^+^) among lymphocytes in the blood, spleen, lung, liver, and bone marrow (BM) from *Ythdf2*^WT^ and *Ythdf2*^ΔNK^ mice (n = 6 per group). **(B)** *Ythdf2*^WT^ and *Ythdf2*^ΔNK^ mice were i.v. injected with an anti-CD45 antibody to mark immune cells, sacrificed after 2 min, and their BM cells were analyzed. Schematic of assays for evaluating NK cell trafficking *in vivo* (upper). Representative plots (lower left) and quantification (lower right) of CD45^+^ NK cells in the BM from *Ythdf2*^WT^ and *Ythdf2*^ΔNK^ mice are shown (n = 4 per group). **(C)** Representative plots (left) and the quantification (right) of immature (CD11b^-^CD27^+^), intermediate mature (CD11b^+^CD27^+^), and terminal mature (CD11b^+^CD27^-^) stages of NK cells from *Ythdf2*^WT^ and *Ythdf2*^ΔNK^ mice are shown (n = 6 per group). **(D)** Representative plots (left) and the quantification (right) of KLRG1^+^NK cells from *Ythdf2*^WT^ and *Ythdf2*^ΔNK^ mice are shown (n = 6 per group). **(E)** A mixture of 5 × 10^6^ BM cells at a 1:1 ratio from CD45.1 or *Ythdf2*^ΔNK^ CD45.2 mice were co-transferred into *Rag2*^−/−^*Il2rg*^−/−^ mice. Reconstitution of recipients was assessed by flow cytometry 8 weeks after transplantation. A schematic diagram for assays evaluating NK cell maturation in a chimeric model (left). The percentages (right) of immature (CD11b^-^CD27^+^), intermediate mature (CD11b^+^CD27^+^), and terminal mature (CD11b^+^CD27^-^) stages of NK cells from CD45.1 mice and *Ythdf2*^ΔNK^ mice are shown (n = 4 per group). Data are shown as mean ± SD and were analyzed by unpaired two-tailed *t*-test (**A, B**), two-way ANOVA with Sidak post-test (**C, D, E**). \**P* < 0.05, \*\**P* < 0.01, \*\*\**P* < 0.001, \*\*\*\**p* < 0.0001. Data are representative of at least three independent experiments.

Immune cells undergo homeostatic proliferation during lymphopenia induced by certain viral infections or caused by chemotherapy (Sun et al., 2011). Although we found YTHDF2 is dispensable for NK cell proliferation at a steady state, we observed a significant decrease in cell proliferation during MCMV infection (Fig. 3, E-G). We, therefore, investigated the role of YTHDF2 in regulating NK cell homeostatic proliferation in a lymphopenic setting *in vivo*. We co-transferred an equal number of splenic NK cells from CD45.2 *Ythdf2*^ΔNK^ mice or CD45.1 congenic mice into lymphocyte-deficient *Rag2*^−/−^*Il2rg*^−/−^ mice. The results showed a greater proportion of NK cells were derived from CD45.1 WT control mice than from CD45.2 *Ythdf2*^ΔNK^ mice day 3 after cell transfer (Fig. S3 D). Further analysis demonstrated that the reduction of NK cells from *Ythdf2*^ΔNK^ mice was due to impaired cell proliferation (Fig. S3 E) but not cell apoptosis (Fig. S3 F), suggesting that YTHDF2 drives NK cell homeostatic proliferation *in vivo* under lymphopenic conditions.

Further differentiation of murine NK cells can be classified into immature (CD11b^-^CD27^+^), intermediate mature (CD11b^+^CD27^+^), and terminal mature (CD11b^+^CD27^-^) stages based on CD11b and CD27 levels (Chiossone et al., 2009; Geiger and Sun, 2016). We found that *Ythdf2* expression increased with maturation and CD11b^-^CD27^+^, CD11b^+^CD27^+^, CD11b^+^CD27^-^ and display lowest, intermediate, and highest expression levels of *Ythdf2*, respectively (Fig. S3 G), indicating that YTHDF2 may be involved in NK cell maturation. We, therefore, investigated the role of YTHDF2 in NK cell maturation defined by the cell surface markers CD11b and CD27. We found that loss of *Ythdf2* in NK cells resulted in a significant decrease in the frequency of terminal mature NK cells and/or an increase in immature and intermediate mature NK cells in the spleen, liver, lung, and blood but not BM (Fig. 4 C and Fig. S3 H), indicating that YTHDF2 positively regulates terminal NK cell maturation. Consistent with these data, the levels of KLRG1, which is a terminal NK cell maturation marker, were significantly lower in *Ythdf2*^ΔNK^ mice in the spleen, liver, and lung but not BM compared to that of *Ythdf2*^WT^ mice in the corresponding organs or tissue compartments (Fig. 4 D). To determine whether the decreased number of mature NK cells by *Ythdf2* deficiency is cell-intrinsic, we created chimeras in *Rag2*^−/−^*Il2rg*^−/−^ mice by injecting BM cells from CD45.1 WT and CD45.2 *Ythdf2*^ΔNK^ mice, mixed at a 1:1 ratio. As shown by flow cytometry at 8 weeks post-transplantation, a reduced proportion of terminal mature NK cells was derived from CD45.2 *Ythdf2*^ΔNK^ BM cells than those from CD45.1 WT control cells (Fig. 4 E), suggesting that NK cell terminal maturation controlled by YTHDF2 is cell intrinsic. T-box transcription factors Eomes and Tbet are critical for NK cell maturation (Daussy et al., 2014; Gordon et al., 2012). Intracellular staining revealed a significant reduction in the protein levels of Eomes in NK cells from *Ythdf2*^ΔNK^ mice compared with *Ythdf2*^WT^ mice (Fig. S3 I). In addition, we found that the reduction of protein and mRNA levels of Eomes specifically occurred in terminal mature (CD11b^+^CD27^-^) NK cells (Fig. S3, J-L). However, in contrast to Eomes, the expression of Tbet was equivalent in NK cells between *Ythdf2*^ΔNK^ and *Ythdf2*^WT^ mice (Fig. S3, I and K), indicating that YTHDF2 possibly regulates NK cells terminal maturation by targeting Eomes.

### YTHDF2 promotes NK Cell effector function

Our finding that *Ythdf2* deficiency impeded NK cell homeostasis and maturation motivated us to determine if YTHDF2 also repressed NK cell effector function. The major mechanism that regulates NK cell function is the relative contribution of inhibitory and activating receptors (Chan et al., 2014). We then checked the expression levels of molecules associated with NK cell activation or inhibition. We found that the expression levels of the activating receptors CD226 and NKG2D, but not CD69, were decreased in splenic NK cells from *Ythdf2*^ΔNK^ mice compared with those of *Ythdf2*^WT^ mice (Fig. 5 A), whereas the inhibitory receptors NKG2A, TIGIT, and PD-1 were similar between *Ythdf2*^ΔNK^ mice and *Ythdf2*^WT^ mice (Fig. 5 A and Fig. S3 M). In addition, *Ythdf2*^ΔNK^ NK cells produced significantly less IFN-γ than *Ythdf2*^WT^ NK cells when stimulated with IL-12 plus IL-18, or with YAC-1 mouse lymphoma cells (Fig. 5, B and C). We then tested the cytotoxic ability of *Ythdf2*^ΔNK^ NK cells and found that *Ythdf2*^ΔNK^ NK cells had significantly reduced cytotoxicity against MHC class I-deficient mouse lymphoma cell lines RMA-S (Fig. 5 D) but showed similar cytotoxicity against MHC class I-sufficient RMA cells (Fig. S3 N) as shown by a ^51^Cr assay. Taken together, these findings demonstrate that YTHDF2 is essential for NK cell effector function.

**Figure 5.**
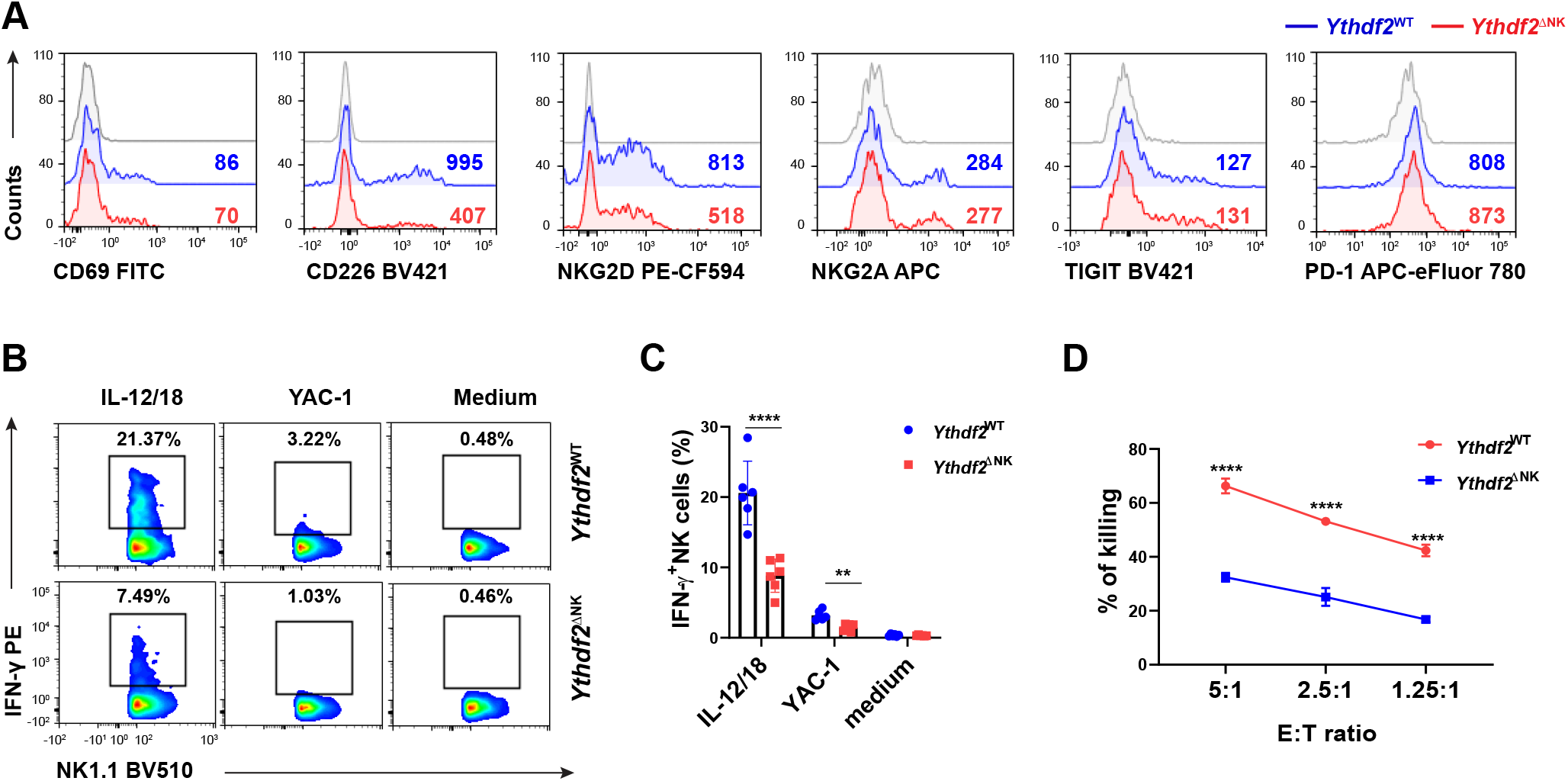
NK cell effector functions require YTHDF2. **(A)** Representative histograms showing expression of the indicated molecules in splenic NK cells from *Ythdf2*^WT^ and *Ythdf2*^ΔNK^ mice. **(B-C)** Representative plots (**B**) and quantification (**C**) of IFN-γ expression by splenic NK cells from *Ythdf2*^WT^ and *Ythdf2*^ΔNK^ mice when stimulated with or without IL-12 plus IL-18 or YAC-1 mouse lymphoma cells (n = 6 per group; at least three independent experiments). **(D)** Mice were treated with an i.p. injection of Poly I:C (200 μg/mice) for 18h. Poly I:C-activated NK cells were isolated from the spleen and co-cultured with MHC class I-deficient RMAS cells at a ratio of 5:1, 2.5:1, and 1.25:1. Cytotoxicity of NK cells was evaluated by standard ^51^Cr release assays (n = 3 per group; at least two independent experiments). Data are shown as mean ± SD and were analyzed by two-way ANOVA with Sidak post-test. \*\**P* < 0.01, \*\*\**P* < 0.001.

### YTHDF2 is required for IL-15-mediated NK cell survival, proliferation, and effector functions

IL-15 is one of, if not the most important cytokine for pleiotropic functions of NK cells. We and others previously discovered that IL-15 plays key roles in regulating NK cell homeostasis, survival, and effector functions (Becknell and Caligiuri, 2005; Carson et al., 1997; Carson et al., 1994; Wang et al., 2019b; Yu et al., 2013). In the current study, we showed that IL-15 upregulated mRNA and protein levels of YTHDF2 in NK cells (Fig. 6A and Fig. 1, D-F). Therefore, we speculated that YTHDF2 is required for IL-15-mediated survival and effector functions of NK cells. We first attempted to identify potential transcription factors downstream of IL-15 signaling that directly regulate YTHDF2 expression. We analyzed the ENCODE using the University of California Santa Cruz (UCSC) Genome Browser Database (https://genome.ucsc.edu), which provides predictions of binding sites across the entire genome, in combination with JASPAR (http://jaspar.genereg.net). We found that STAT5, which is a key downstream factor of IL-15 in NK cells, has four binding sites within 2 Kb upstream of the transcription start site (TSS) of *Ythdf2*, indicating that IL-15 may positively regulate *Ythdf2* transcription in mouse NK cells through STAT5. By utilizing STAT5 inhibitor, STAT5-IN-1 (Muller et al., 2008), we showed that inhibition of STAT5 resulted in a decrease of *Ythdf2* at both mRNA and protein levels in murine NK cells (Fig. S4, A and B). To further confirm YTHDF2 is a downstream factor regulated by STAT5, we used *Stat5^fl/fl^ Ncr1-iCre* mice (hereafter referred to as *Stat5*^ΔNK^ mice), where STAT5 is specifically deleted in mouse NK cells (Wiedemann et al., 2020). We treated splenic NK cells from *Stat5*^WT^ and *Stat5*^ΔNK^ mice with IL-15. We found that YTHDF2 substantially decreased in NK cells from *Stat5*^ΔNK^ mice compared to NK cells from *Stat5*^WT^ mice at both the mRNA and protein levels (Fig. 6, B and C; and Fig. S4, C and D). Luciferase reporter assay showed that both STAT5a and STAT5b activated *Ythdf2* gene transcription directly (Fig. 6, D and E). Chromatin immunoprecipitation (ChIP)- qPCR results showed that STAT5 has a significant enrichment on four sites over normal IgG control, indicating a direct binding in mouse NK cells (Fig. 6 F). Together, our results demonstrate that YTHDF2 expression is regulated by STAT5 downstream of IL-15 signaling in NK cells.

**Figure 6.**
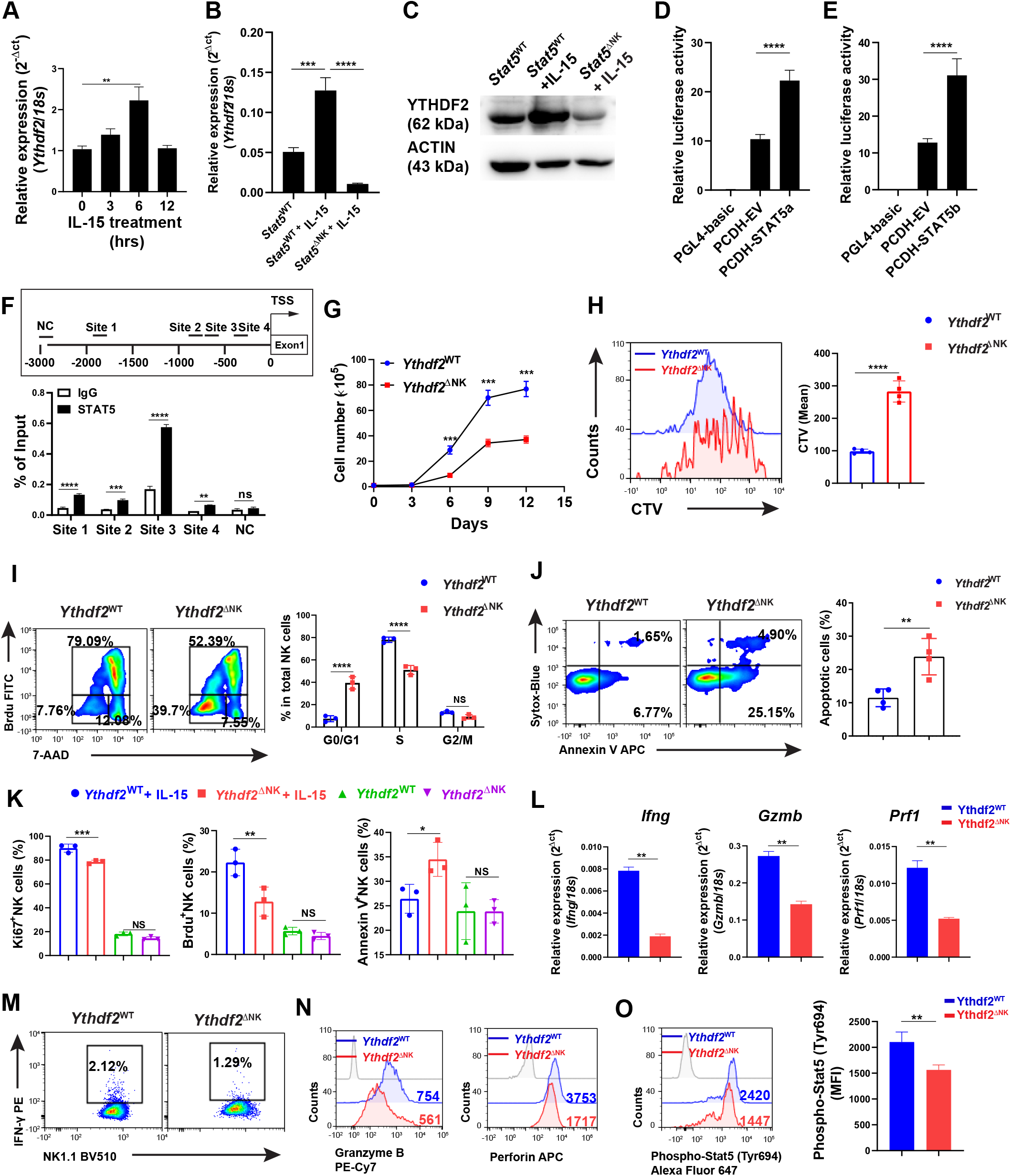
YTHDF2 contributes to IL-15-mediated NK cell survival, proliferation, and effector functions. **(A)** qPCR showing the expression of *Ythdf2* in NK cells at indicated time points following stimulation of IL-15**. (B)** qPCR showing the expression of *Ythdf2* in NK cells from *Stat5*^WT^ and *Stat2*^ΔNK^ mice under the stimulation of IL-15. **(C)** Immunoblotting showing the YTHDF2 protein levels in splenic NK cells from *Stat5*^WT^ and *Stat5*^ΔNK^ mice under the stimulation of IL-15. **(D-E)** Luciferase reporter assay shows that both STAT5a **(D)** and STAT5b **(E)** activate *Ythdf2* gene transcription. **(F)** Scheme denoting putative STAT5 binding sites in the *Ythdf2* promoter (upper). Binding of STAT5 to the *Ythdf2* promoter in NK cells as determined by chromatin ChIP– qPCR (lower). **(G)** Splenic NK cells isolated from *Ythdf2*^WT^ and *Ythdf2*^ΔNK^ were cultured *in vitro* in the presence of IL-15 (50 ng/ml), followed by enumeration by a trypan blue exclusion assay. **(H)** Representative histograms (left) and quantification (right) of CTV dilution of NK cells from *Ythdf2*^WT^ and *Ythdf2*^ΔNK^ mice five days after *in vitro* culture with IL-15 (50 ng/ml). **(I)** Representative plots (left) and quantification (right) of cell cycle distribution of NK cells from *Ythdf2*^WT^ and *Ythdf2*^ΔNK^ mice five days after *in vitro* culture with IL-15 (50 ng/ml). **(J)** Representative plots (left) and quantification (right) of apoptotic (Annexin V^+^Sytox-Blue^+/-^) NK cells from *Ythdf2*^WT^ and *Ythdf2*^ΔNK^ mice five days after *in vitro* culture with IL-15 (50 ng/ml). (**K**) Mice were treated with IL-15 (2 μg per day) for 5 days (n = 3 per group). On day 4, mice were injected i.p. with Brdu (5 mg per mice) overnight. Splenic NK cells were then isolated, followed by Ki67 cell proliferation and Annexin V apoptosis assessment, measured by flow cytometry. **(L)** qPCR showing the expression of *Ifng, Gzmb*, and *Prf1* in NK cells from *Ythdf2*^WT^ and *Ythdf2*^ΔNK^ mice following stimulation of IL-15 (50 ng/ml) overnight. **(M-N)** Representative plots and histograms of IFN-γ (**M**), granzyme B (**N**), and perforin (**N**) levels in NK cells from *Ythdf2*^WT^ and *Ythdf2*^ΔNK^ mice following stimulation with IL-15 (50 ng/ml) overnight. **(O)** Representative histograms (left) and quantification (right) of pSTAT5 in NK cells from *Ythdf2*^WT^ and *Ythdf2*^ΔNK^ mice after *in vitro* stimulation of IL-15 (50 ng/ml) for 1 h. Data are shown as mean ± SD and were analyzed by unpaired two-tailed *t*-test (**H, J, L, O**), one-way ANOVA (**A, B, D, E, K**), or two-way ANOVA with Sidak post-test (**F, G, I**). \**P* < 0.05, \*\**P* < 0.01, \*\*\**P* < 0.001, \*\*\*\**P* < 0.0001. Data are representative of at least two independent experiments.

We then wondered whether *Ythdf2*-deficient NK cells were defective in their response to IL-15. We isolated splenic NK cells from *Ythdf2*^ΔNK^, mice and *Ythdf2*^WT^ mice and cultured *in vitro* in the presence of IL-15. We found that NK cell growth was significantly decreased in *Ythdf2*^ΔNK^ mice compared to *Ythdf2*^WT^ mice (Fig. 6 G). CellTrace Violet labeling assay showed that the proliferation of NK cells was impaired by *Ythdf2* deficiency (Fig. 6 H). Cell cycle distribution analysis revealed a significantly increased fraction of NK cells from *Ythdf2*^ΔNK^ mice in G_0_/G_1_ phase but a significantly decreased fraction of NK cells from *Ythdf2*^ΔNK^ mice in S phase (Fig. 6 I), suggesting *Ythdf2* deficiency results in G_0_/G_1_ phase arrest in NK cells. In addition, there was a two- to three-fold increase in apoptotic (Annexin V^+^Sytox-Blue^+/-^) NK cells from *Ythdf2*^ΔNK^ mice compared to those from *Ythdf2*^WT^ mice (Fig. 6 J), indicating the defective survival of NK cells after the loss of *Ythdf2*. These results suggest that YTHDF2 regulates the responsiveness of NK cells to IL-15 *in vitro*. Since IL-15 is poorly translated and secreted *in vivo* at a steady-state (Corbel et al., 1996; Fehniger et al., 2001), to explore whether *Ythdf2*-deficient NK cells are defective in their response to IL-15 *in vivo*, we treated mice with IL-15. We found that *Ythdf2*-deficient vs. WT NK cells showed significantly reduced cell proliferation and increased cell apoptosis when mice were treated with IL-15 (Fig. 6 K and Fig. S4, E-J). These data suggest that YTHDF2 regulates the responsiveness of NK cells to IL-15, especially under some conditions with a high level of IL-15. We also found that the mRNA and protein levels of IFN-γ, granzyme B, and perforin were significantly reduced in *Ythdf2*^ΔNK^ NK cells compared with *Ythdf2*^WT^ NK cells in response to IL-15 (Fig. 6, L-N), indicating that YTHDF2 also contributes to IL-15-mediated NK cell effector functions in *vitro*. Collectively, our data demonstrate that YTHDF2 is required for IL-15-mediated NK cell survival, proliferation, and effector functions.

Next, we investigated the downstream mechanisms by which YTHDF2 regulates IL-15-mediated NK survival, proliferation, and effector function. We found that NK cells from *Ythdf2*^ΔNK^ mice and *Ythdf2*^WT^ mice had similar levels of the IL-15 receptors CD122 (IL-15Rβ) and CD132 (IL-15Rγc) (Fig. S4 K). IL-15 signaling is mediated by at least three downstream signaling pathways in NK cells: Ras–Raf–MEK–ERK, PI3K–AKT–mTOR, and JAK1/3– STAT3/5 (Mishra et al., 2014). To investigate which signaling pathway is regulated by YTHDF2 in NK cells, we examined the phosphorylation levels of ERK, AKT, S6, STAT3, and STAT5 in NK cells from *Ythdf2*^ΔNK^ mice or *Ythdf2*^WT^ mice after stimulation with IL-15. The results showed that *Ythdf2* deficiency did not affect ERK, AKT, S6, and STAT3 phosphorylation upon IL-15 stimulation (Fig. S4, L and M), but significantly inhibited STAT5 activation, as evidenced by reduced phosphorylation levels of STAT5 in *Ythdf2*-deficient NK cells than that in NK cells from *Ythdf2*^WT^ mice (Fig. 6 O), indicating that YTHDF2 is required for the optimum IL-15/STAT5 signaling in activated NK cells. Because we showed that phosphorylated STAT5 downstream of IL-15 binds to the promoter of *Ythdf2* (Fig. 6, D-F), and here we demonstrated that YTHDF2 is required for optimum STAT5 phosphorylation, our data suggest a STAT5-YTHDF2 positive feedback loop downstream of IL-15 that may control NK cell survival, proliferation, and effector functions.

### Transcriptome-wide analysis identifies *Tardbp* as a YTHDF2 target in NK cells

To address the molecular mechanism by which YTHDF2 regulates NK cells, we first performed RNA-seq in IL-2-expanded *Ythdf2*^WT^ and *Ythdf2*^ΔNK^ NK cells. The deletion of *Ythdf2* in NK cells resulted in 617 differentially expressed genes (DEGs), including 252 up-regulated genes and 365 down-regulated genes (Fig. S5 A). Gene Ontology (GO) analysis showed that the DEGs were significantly enriched in cell cycle, cell division, and cell division-related processes, including mitotic cytokinesis, chromosome segregation, spindle, nucleosome, midbody, and chromosome (Fig. 7 A). Gene Set Enrichment Analysis (GSEA) demonstrated significant enrichment of E2F targets, G2/M checkpoint, and mitotic spindle hallmark gene sets in *Ythdf2*^ΔNK^ NK cells (Fig. S5 B). Cell cycle and division-related genes, including *Aurka, Aurkb, Cdc20, Cdc25b, Cdc25c, Cdk1, E2f2*, and *Plk1* (Bertoli et al., 2013), were significantly decreased in NK cells from *Ythdf2*^ΔNK^ mice (Fig. 7 B). Spindle and chromosome segregation genes such as *Anln, Aspm, Birc5, Bublb, Cenpe, Esco2, Ska1, Ska3*, and *Tpx2* (Gorbsky, 2015), were also significantly downregulated in NK cells from *Ythdf2*^ΔNK^ mice compared to NK cells from *Ythdf2*^WT^ mice (Fig. 7 C). In addition, cell survival genes, including *Birc5* (Niu et al., 2010), *Septin4* (Larisch et al., 2000), and *Rffl* (Yang et al., 2007) were significantly decreased in NK cells from *Ythdf2*^ΔNK^ mice (Fig. 7 D). NK cell effector function genes, including *Klrk1, Ncr1, CD226*, and *Gzma* (Bezman et al., 2012), were also significantly downregulated in NK cells from *Ythdf2*^ΔNK^ mice (Fig. 7 D). These data support our characterized roles of YTHDF2 in regulating NK proliferation, survival, and effector functions.

**Figure 7.**
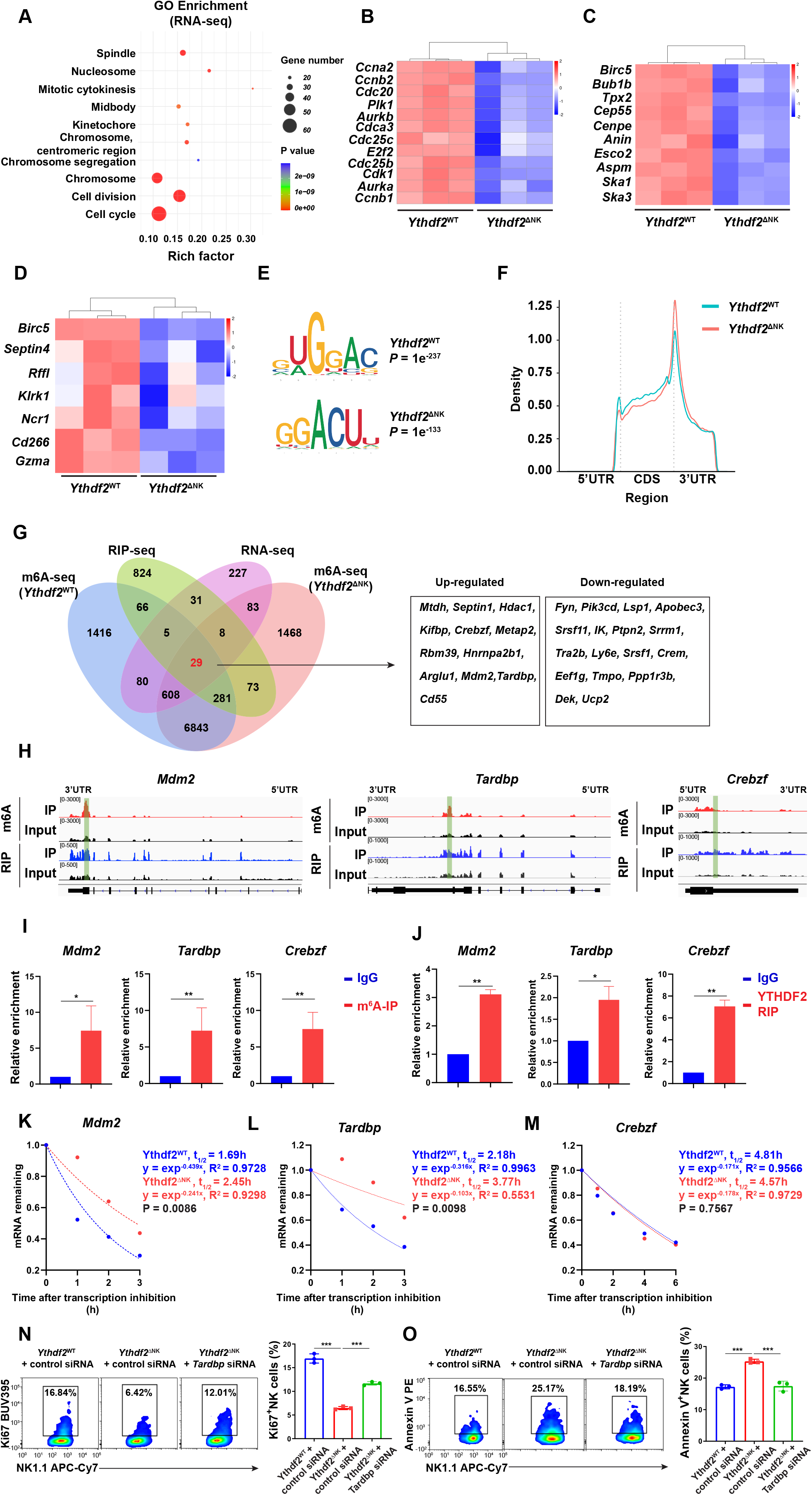
Transcriptome-wide identification of YTHDF2-binding targets in NK cells. **(A)** Top 10 GO clusters from GO analysis of differentially expressed genes from RNA-seq data. **(BD)** Heatmaps of differentially expressed genes between NK cells from *Ythdf2*^WT^ and *Ythdf2*^ΔNK^ mice from RNA-seq grouped by cell cycle and division (**B**), spindle and chromosome (**C**), and cell survival and NK cell function (**D**). **(E)** The m^6^A motif detected by the HOMER motif discovery toll with m^6^A-seq data. **(F)** Density distribution of the m^6^A peaks across mRNA transcriptome from m^6^A-seq data. **(G)** Overlapping analysis of genes identified by RNA-seq, m^6^A-seq, and RIP-seq. Twelve up-regulated and 17 down-regulated differentially expressed transcripts bound by YTHDF2 and marked with m^6^A are listed on the right tables. **(H)** Distribution of m^6^A peaks and YTHDF2-binding peaks across the indicated transcripts by Integrative Genomics Viewer. **(I and J)** RIP using either m^6^A (**I**) or YTHDF2 antibody (**J**) following qPCR to validate the target genes identified by m^6^A-seq and RIP-seq. Rabbit IgG was used as control. Enrichment of indicated genes was normalized to the input level. **(K-M)** The mRNA half-life (t_1/2_) of *Mdm2* (**K**), *Tardbp* (**L**), and *Crebzf* (**M**) transcripts in NK cells from *Ythdf2*^WT^ and *Ythdf2*^ΔNK^ mice. Data represent two independent experiments. (**N-O**) IL-2 expanded NK cells were transfected with *Tardbp* specific siRNA cells under the stimulation of IL-15 (50 ng/ml). Three days post transfection, cell proliferation or apoptosis was analyzed by Ki67 staining or Annexin V straining, respectively, followed by a flow cytometric analysis (n = 3 per group, at least two independent experiments). Data are shown as mean ± SD and were analyzed by unpaired two-tailed *t*-test (**I, J**), one-way ANOVA (**N, O**). \**P* < 0.05, \*\**P* < 0.01, \*\*\**P* < 0.001.

We then performed m^6^A-seq in IL-2 expanded NK cells from *Ythdf2*^WT^ and *Ythdf2*^ΔNK^ mice. Principal component analysis (PCA) showed that three biological replicates of each genotype clustered together (data not shown), suggesting good repeatability of m^6^A-seq samples. HOMER analysis identified the m^6^A consensus motif (GGAC), indicating the successful enrichment of m^6^A modified transcripts (Fig. 7 E). m^6^A modifications were predominantly located in proteincoding transcripts, and the peaks were enriched in the 5’UTR and 3’UTR regions, especially around the start and stop codons (Fig. 7 F and Fig. S5 C). GO enrichment analysis of genes with m^6^A peaks revealed that most m^6^A marked transcripts in NK cells were enriched in pathways involved in cell cycle and cell proliferation (Fig. S5 D).

It is well known that YTHDF2 acts as an m^6^A reader, which recognizes and binds m^6^A-containing RNAs and regulates the degradation of m^6^A transcripts. We then performed RNA immunoprecipitation sequencing (RIP-Seq) using YTHDF2 antibody to map the target transcripts bound by YTHDF2 in NK cells. YTHDF2-binding sites were enriched in the proteincoding sequence and 3’UTR regions (Fig. S5, E and F). We identified 3951 potential YTHDF2 binding peaks from 1290 transcripts, 426 (33%) of which were detected with significant m^6^A enrichment. GO enrichment analysis showed that the YTHDF2 target 1290 transcripts were enriched in protein translation, RNA binding, and RNA splicing (Fig. S5 G).

We then identified the potential target transcripts from overlapping transcripts through the RNA-seq, m^6^A-seq, and RIP-seq and found a set of 29 transcripts bound by YTHDF2, marked with m^6^A (in both *Ythdf2*^WT^ and *Ythdf2*^ΔNK^ NK cells) were differentially expressed between NK cells from *Ythdf2*^WT^ and *Ythdf2*^ΔNK^ mice (Fig. 7 G). Among them, 12 transcripts were upregulated, and 17 transcripts were downregulated in NK cells from *Ythdf2*^ΔNK^ mice compared to those from *Ythdf2*^WT^ NK cells (Fig. 7 G). Based on our functional characterization of YTHDF2 and the GO analysis from the RNA-seq data, *Ythdf2* deficiency affects NK cell division and proliferation, possibly by facilitating mRNA degradation of cell cycle checkpoints or negative regulators during cell division. Among the 12 upregulated transcripts, three of them have been reported to negatively regulate cell division or proliferation, including, *Mdm2* (murine double minute 2, MDM2) (Frum et al., 2009; Giono and Manfredi, 2007; Giono et al., 2017), *Tardbp* (TAR DNA-binding protein 43, TDP-43) (Ayala et al., 2008; Sanna et al., 2020), and *Crebzf* (CREB/ATF BZIP Transcription Factor, CREBZF) (Hu et al., 2020; Lopez-Mateo et al., 2012), suggesting that they are the potential targets of YTHDF2 in NK cells. Of note, the m^6^A peaks fit well with the YTHDF2-binding sites at 3’UTR region of *Mdm2* and *Tardbp*, and with the 5’UTR region of *Crebzf* genes, as shown by Integrative Genomics Viewer (Fig. 7 H). RIP using either m^6^A or YTHDF2 antibody following qPCR confirmed that *Mdm2, Tardbp*, and *Crebzf* were indeed m^6^A methylated and enriched predominately by YTHDF2 in NK cells (Fig. 7, I and J), supporting m^6^A-Seq and RIP-seq data, which indicate that these three genes are potential targets of YTHDF2 in NK cells. To investigate whether YTHDF2 regulates *Mdm2, Tardbp*, and *Crebzf* expression through modulating the mRNA stability, we measured the mRNA degradation of the three targets by inhibition of transcription with actinomycin D in NK cells from *Ythdf2*^ΔNK^ and *Ythdf2*^WT^ mice. The results showed that *Mdm2* and *Tardbp*, but not *Crebzf* had longer half-lives in NK cells from *Ythdf2*^ΔNK^ mice compared with those from *Ythdf2*^WT^ mice (Fig. 7, K and M), suggesting that *Mdm2* and *Tardbp* are directly regulated by YTHDF2 in NK cells. Using immunoblotting, we confirmed that the protein levels of MDM2 and TDP-43 were upregulated in NK cells from *Ythdf2*^ΔNK^ mice (Fig. S5 H). To further confirm these two genes are functional targets of YTHDF2, we used *Mdm2* or *Tardbp* specific siRNA to knock down the expression of the two genes *in vitro* (Fig. S5, I and J). We then compared cell proliferation and survival of *Ythdf2*^ΔNK^ NK cells with vs. without knockdown of *Mdm2* or *Tardbp* in the presence of IL-15. The results showed that knock-down of *Tardbp* could at least partially rescue the defect in cell proliferation and cell survival in NK cells from *Ythdf2*^ΔNK^ mice (Fig. 7, N and O). However, knock-down of *Mdm2* had no rescue effects (Fig. S5, K and L). These results indicate that YTHDF2 regulates NK cell proliferation and division at least partially through inhibiting the mRNA stability of *Tardbp*.

## Discussion

In this study, we reported the multifaceted roles of YTHDF2-mediated m^6^A methylation in NK cell immunity. We found that YTHDF2, one of the most important readers of m^6^A modifications, was critical for maintaining NK cell homeostasis, maturation, IL-15-mediated survival, as well as anti-tumor and anti-viral activity. We also identified a novel positive feedback loop between STAT5 and YTHDF2, downstream of IL-15 that contributes to effector functions and survival in mouse NK cells. Our study elucidates the biological roles of YTHDF2 or m^6^A methylations in general in NK cell innate immunity. It fills in the gap of knowledge as to how YTHDF2 regulates the innate immune response to malignant transformation and viral infection. Our findings provide a new direction to harness the NK cell anti-tumor immunity and simultaneously advance our understanding of the m^6^A modifications in shaping innate immunity.

The m^6^A reader protein YTHDF2 regulates the stability of target mRNAs (Du et al., 2016; Wang et al., 2014). Numerous studies have supported the broad impact of YTHDF2 in various biological processes. YTHDF2 suppresses normal hematopoietic stem cell (HSC) expansion and self-renewal but is also required for long-term HSC maintenance (Li et al., 2018; Mapperley et al., 2020; Paris et al., 2019; Wang et al., 2018a). Recently, this gene has been found to be involved in restraining inflammation during bacterial infection (Wu et al., 2020a). The role and mechanism of YTHDF2 in tumor development have been well-studied. YTHDF2 promotes leukemic stem cell development and acute myeloid leukemia (AML) initiation (Paris et al., 2019). Besides leukemia, YTHDF2 has been shown to promote the development of solid tumors, including prostate cancer (Li et al., 2020), glioblastoma (Dixit et al., 2020), and hepatocellular carcinoma (Chen et al., 2018; Hou et al., 2019; Zhang et al., 2020) by targeting diverse m^6^A-modified transcripts, such as tumor suppressors LHPP and NKX3-1 (Li et al., 2020), IGFBP3 (Dixit et al., 2020), OCT4 (Zhang et al., 2020), and SOCS2 (Chen et al., 2018). Although YTHDF2 plays a promoting role in tumor progression, our study reveals a beneficial role of YTHDF2 in the immune response to tumor cells, particularly in NK cells, which is a key component of innate immunity against viral infections and malignant transformation. Therefore, future development of YTHDF2 inhibitors to target tumor cells for cancer therapy should be pursued with caution as inhibition of YTHDF2 may impair the host anti-tumor responses by NK cells. On the other hand, harnessing anti-tumor activity of YTHDF2 in NK or other immune cells should consider its potential effect acting on tumor cells directly. Differential targeting of YTHDF2 in tumor cells and in immune cells should maximize anti-tumor activity.

NK cells mainly utilize two main approaches to destroy tumor cells and virus-infected cells: 1) release of cytotoxic molecules such as perforin and granzymes that directly induce target cell apoptosis or pyroptosis (Zhou et al., 2020); 2) secretion of several cytokines such as IFN-γ, TNF-α, granulocyte macrophage colony-stimulating factor (GM-CSF), and chemokines (MIP-1α, MIP-1β, IL-8, and RANTES) that enhance the function of other innate and adaptive immune cells (Fauriat et al., 2010; Reiter, 1993). In this study, we observed non-identical mechanisms by which YTHDF2 regulates NK cell anti-tumor and anti-viral immunity. In the tumor setting, YTHDF2 promotes the secretion of perforin, granzyme B, and IFN-γ by NK cells for controlling melanoma metastasis, whereas during MCMV infection, YTHDF2 promotes NK cell-mediated anti-viral activity against MCMV mainly through regulating perforin. One potential explanation for this discrepancy is that NK cell activation by tumor or virus is regulated in different manners. MCMV activates NK cells by encoding protein m157, a ligand of NK cell receptor Ly49H (Smith et al., 2002), and associates with two intracellular adaptors, DAP10 and DAP12 (French et al., 2006; Orr et al., 2009). However, in the tumor context, NK cell activation is tightly regulated by its interaction with different NK cell receptor ligands expressed by tumor cells, as well as the cytokines such as IL-15 and TGF-β in the tumor microenvironment (Wu et al., 2020b). In addition, we found that YTHDF2 promotes NK cell maturation by regulating Eomes. However, no binding sites were found in Eomes mRNA, suggesting that YTHDF2 indirectly regulates Eomes expression. Further studies are warranted to investigate the mechanism by which YTHDF2 regulates Eomes expression. In addition, how the YTHDF2 receives and mediates signals can quickly shape the NK cell immune response against virus or tumor cells differently also requires further exploration.

Adoptive transfer of allogeneic NK cells into leukemia patients can lead to remission (Ruggeri et al., 2002; Yilmaz et al., 2020). Chimeric antigen receptor (CAR)- engineered NK cells have been shown to provide significant benefits in relapsed or refractory CD 19-positive lymphoma and leukemia (Liu et al., 2020). However, limited expansion and persistence of NK cells *in vivo*, as well as limited NK cell trafficking and infiltration into tumor sites remain a major challenge for NK cell-based therapy (Yilmaz et al., 2020). Since cancer patients or virally-infected patients, such as those with COVID-19, usually undergo transient lymphopenia (Grossman et al., 2015; Zhao et al., 2020), efficient expansion of NK cells during lymphopenia is critical for controlling tumor growth and viral infection. Our findings show that YTHDF2 drives NK cell egress from BM and promotes NK cell homeostatic proliferation during lymphopenia *in vivo* in mice lacking T, B, and NK cells. Our study also shows that YTHDF2 positively regulates NK cell effector function. Therefore, incorporation of YTHDF2 expression into NK or CAR-NK cells may have multifaceted benefits for NK cell expansion during manufacturing *in vitro*, persistence, and enhancement of effector function *in vivo*. Furthermore, the upregulation of YTHDF2 that we observed in the tumor setting and during viral infection may increase the ability of NK or CAR-NK cells to infiltrate into the disease microenvironment.

IL-15 is a key regulator of NK cell development, homeostasis, survival, and effector function (Becknell and Caligiuri, 2005; Mishra et al., 2014; Yu et al., 2013). Our group previously reported a novel IL-15-AKT-XBP1s signaling pathway that contributes to the effector functions and survival of human NK cells (Wang et al., 2019b). However, the exact mechanism(s) by which IL-15 regulates NK cell survival has not yet been fully understood. Here we found another novel mechanism in that STAT5-YTHDF2 forms a positive feedback loop downstream of IL-15 in mouse NK cells that in turn controls NK cell proliferation, survival, and effector functions. Our previous report showed that IL-15 does not induce transcription of XBP1s, and XBP1s does not interact with STAT5 in NK cells (Wang et al., 2019b), suggesting that regulation of XBP1s by IL-15 is STAT5-independent. We therefore identify two novel mediators of IL-15 in NK cells, XBP1s and YTHDF2, for which XBP1s is not regulated by STAT5, while YTHDF2 is STAT5-dependent. This complexity of characterized IL-15 signaling may match the complex and pleiotropic role of IL-15, which is a key component of both the inflammatory milieu in the tumor microenvironment and the response to viral infection (Nguyen et al., 2002; Santana Carrero et al., 2019). The complexity is also reflected in that *Ythdf2* deficiency has no effect on the survival and proliferation of resting NK cells *in vivo*, while YTHDF2 plays a critical role in regulating proliferation and/or survival of NK cells activated by IL-15 or by MCMV infection. Our study also supports a concept that YTHDF2 or m^6^A in general plays a more central role in NK cell dynamics in the activated state and/or disease settings.

In this study, we applied a multi-omics strategy (RNA-seq, m^6^A-seq, and RIP-seq) to identify the targets of YTHDF2 in NK cells. In line with our finding that *Ythdf2*-deficient NK cells showed significantly delayed cell growth, we found a large number of genes related to cell cycle and cell division that were markedly decreased in *Ythdf2*^ΔNK^ NK cells, suggesting YTHDF2 controls cell growth by regulating cell cycle. Of note, the m^6^A modifications have been widely involved in regulating cell cycle. METTL3 promotes cell growth of AML by enhancing the translation of genes in the cell cycle pathway (Barbieri et al., 2017; Vu et al., 2017). METTL14 deletion extends cortical neurogenesis into postnatal stages by prolonging the S to M phase transition of radial glia cells (Yoon et al., 2017). Consistent with our study, it was reported that in HeLa cells, YTHDF2 targets pathways not only involved in molecular function but also in cell proliferation and survival (Wang et al., 2014); Fei et al. recently reported that YTHDF2 promoted cell proliferation possibly by facilitating mRNA degradation during cell cycle in HeLa cells (Fei et al., 2020), suggestive of a universal mechanism(s) of YTHDF2 or its associated m^6^A modifications in maintaining cell survival and function. Because millions of cells are needed for the multi-omics strategy analysis, we had to rely on *ex vivo* expanded and highly proliferative NK cells. This can explain the observation that the screened targets of YTHDF2 are mainly cell-cycle genes, while potential targets that control cell survival, effector function, and maturation of NK cells were not shown. Regardless, we believe our data provide the first evidence that YTHDF2 targets contribute to cell cycle and cell division processes in immune cells, particularly in NK cells.

In conclusion, we discovered a previously unknown role of YTHDF2 or m^6^A methylation as a positive regulator of NK cell anti-tumor and anti-viral activity, as well as NK cell homeostasis and maturation. These findings provide insight into how NK cells effectively survey against tumor metastases and viral infection through m^6^A mRNA methylation.

## Materials and methods

### Mice

*Ythdf2*^fl/fl^ mice were generated by the lab of Jianjun Chen (City of Hope, USA). *Ncrl*-iCre mice were a gift of Eric Vivier (Centre d’Immunologie de Marseille-Luminy, Marseille, France) (Narni-Mancinelli et al., 2011). *Rag2*^−/−^*Il2rg*^−/−^ mice were a gift of Flavia Pichiorri (City of Hope, USA). NCI B6-Ly5.1/Cr mice (CD45.1 mice) were purchased from Charles River Laboratories. IL-15 transgenic (IL-15Tg) mice were generated by our group and backcrossed to C57BL/6 background (Fehniger et al., 2001). *Stat5*^fl/fl^ mice were originally from John J. O’Shea (National Institute of Arthritis and Musculoskeletal and Skin Diseases, USA)(Cui et al., 2004). *Stat5*^fl/fl^ *Ncr1*-iCre mice were generated in the lab of Joseph C. Sun (Memorial Sloan Kettering Cancer Center, New York, USA) (Wiedemann et al., 2020). All mice were on a C57BL/6 background for > 10 generations. Six to twelve-week-old male and female mice were used for the experiments. Cre negative littermates were used as WT controls. All animal experiments were approved by the City of Hope Institutional Animal Care and Use Committee.

### Metastatic melanoma model and MCMV challenge

B16F10 cells (1 × 10^5^) were injected intravenously (i.v.) into mice. Fourteen days after injection, the mice were euthanized for post-mortem analysis. Metastases nodules in the lung were analyzed by macroscopic and counted. The B16F10 cell line was provided by Hua Yu (City of Hope, USA). *Ythdf2*^ΔNK^ and *Ythdf2*^WT^ mice were infected with intraperitoneal (i.p.) injection of Smith strain MCMV (2.5 × 10^4^ PFU), which was purchased from American Type Culture Collection (VR-1399; Manassas, VA). Peripheral blood samples were obtained through submandibular puncture on days 0, 4, and 7 days after infection. To measure viral loads in the peripheral blood, spleen, and liver, DNA was isolated using a QIAGEN DNeasy Blood and Tissue Kit for qPCR analysis. The following primers were used: MCMV-*IE1*, 5’-AGCCACCAACATTGACCACGCAC-3’ (forward) and MCMV-*IE1*, 5’-GCCCCAACCAGGACACACAACTC-3’ (reverse).

### *In vivo* mouse treatment with IL-15

For *in vivo* mouse treatment with IL-15, *Ythdf2*^ΔNK^ and *Ythdf2*^WT^ mice were i.p. injected with 2 μg recombinant human IL-15 (National Cancer Institute, Cat No. 745101) for five days. The mice were then euthanized for a flow cytometric analysis.

### Flow cytometry

Single-cell suspensions were prepared from the bone marrow, blood, spleen, liver, and lung of *Ythdf2*^ΔNK^ and *Ythdf2*^WT^ mice as described previously (Wang et al., 2018b). Flow cytometry analysis and cell sorting were performed on BD LSRFortessa X-20 and FACSAria Fusion Flow Cytometer (BD Biosciences), respectively. Data were analyzed using NovoExpress Software (Agilent Technologies). The following fluorescence dye–labeled antibodies from BD Biosciences, BioLegend, Invitrogen, or Cell Signaling Technology were used: CD3ε (145-2C11), CD19 (1D3), Gr-1 (RB6-8C5), TER-119 (TER-119), CD11c (N418), CD4 (GK1.5), CD8 (53-6.7), CD122 (5H4), CD132 (TUGm2), NK1.1(PK136,), CD11b (M1/70), CD27 (LG.3A10), CD117 (2B8), CD127 (SB/199), CD135 (A2F10.1), KLRG1 (2F1), CD45 (30-F11), CD45.1 (A20), CD45.2 (104), IFN-γ (XMG1.2), 2B4 (m2B4), Granzyme B (QA16A02), Perforin (S16009A), Ly49H (3D10), Ly49D (4e5), CD69 (H1.2F3), CD226 (TX42.1), NKG2D (CX5), NKG2A (16A11), NKp46 (29A1.4), TIGIT (1G9), PD-1 (J43), Annexin V, Ki67 (b56), Tbet (4b10), and Emoes (WD1928), Phospho-p44/42 Erk1/2 (Thr202/Tyr204) (#9101), Phospho-Akt (Ser473) (#5315), Phospho-S6 Ribosomal Protein, Phospho-Stat3 (Tyr705) (#9145), and Phospho-Stat5 (Tyr694) (#9539). For the evaluation of NK cell proliferation, cells were labeled with 5 μM CellTrace Violet (Invitrogen) according to the manufacturer’s protocol before transferring them into recipient mice. Intracellular staining of Ki-67, Tbet, and Eomes was performed by fixing and permeabilizing with the Foxp3/Transcription Factor Staining Kit (eBioscience). For detection of phosphorylated proteins, purified splenic NK cells were pretreated with recombinant human IL-15 (50 ng/ml) for 1h and then fixed with BD Phosflow Fix Buffer I, followed by permeabilization with Phosflow Perm buffer III (BD Biosciences) and staining with antibodies.

### Adoptive cell transfer

For assessing the effect of *Ythdf2* deficiency on NK cell maturation, a mixture of 5 × 10^6^ BM cells at a 1:1 ratio from CD45.1 or *Ythdf2*^ΔNK^ CD45.2 mice were co-transferred into *Rag2*^−/−^*Il2rg*^−/−^ mice. Reconstitution of recipients was assessed by flow cytometry 8 weeks after transplantation. For lymphopenia-induced homeostatic proliferation experiments, equal numbers of purified splenic NK cells from CD45.1 or *Ythdf2*^ΔNK^ CD45.2 mice were co-transferred into *Rag2*^−/−^*Il2rg*^−/−^ mice, followed by assessment of the relative percentages of transferred WT and *Ythdf2*^ΔNK^ NK cells in the spleen of *Rag2*^−/−^*Il2rg*^−/−^ recipients by flow cytometry at indicated time points. For the metastatic melanoma model, 1 × 10^6^ IL-2 expanded NK cells from *Ythdf2*^ΔNK^ or *Ythdf2*^WT^ mice were i.v. injected into *Rag2*^−/−^*Il2rg*^−/−^ mice. One day later, B16F10 cells (1 × 10^5^) were i.v. injected into mice. Fourteen days after injection, mice were euthanized for post-mortem analysis. In some experiments, cells were labeled with CTV (5 μM, Invitrogen) to trace cell proliferation before transfer.

### *In vivo* trafficking assay

For detecting NK cell trafficking from BM to peripheral blood, 1 μg of APC-labeled anti-CD45 antibody was i.v. injected into C57BL/6 mice. Two minutes after the antibody injection, the mice were euthanized immediately, and the BM cells were collected for flow cytometry after the cells were stained with CD3 and NK1.1 antibodies. Parenchymal NK cells were identified by lack of CD45 staining, whereas sinusoidal NK cells were identified by the presence of CD45 labeling. Therefore, the ratio of NK cells in the sinusoids (CD45^+^) to that in the parenchymal regions (CD45^-^) indicates the NK cell trafficking from BM to peripheral blood under a steady state.

### Quantitative real-time RT-PCR (qPCR) and Immunoblotting

RNA was isolated using an RNeasy Mini Kit (QIAGEN) and then reverse transcribed to cDNA with PrimeScript RT Reagent Kit with gDNA Eraser (Takara Bio) following the instructions provided by the manufacturer. mRNA expression levels were analyzed using SYBR Green PCR Master Mix and a QuantStudio 12K Flex Real-Time PCR System (both from Thermo Fisher Scientific). Primer sequences are listed in Supplemental Table 1. Immunoblotting was performed according to standard procedures, as previously described (Deng et al., 2015; Yu et al., 2006). The following antibodies were used: METTL3 (Proteintech, Cat No. 15073-1-AP), METTL14 (Proteintech, Cat No. 26158-1-AP), YTHDF1 (Proteintech, Cat No. 17479-1-AP), YTHDF2 (MBL, Cat No. RN123PW), YTHDF3 (Proteintech, Cat No. 25537-1-AP), ALKBH5 (Abcam, Cat No. ab195377), FTO (Abcam, Cat No. ab124892), MDM2 (Invitrogen, Cat No. 33-7100), TDP-43 (Invitrogen, Cat No. PA5-29949), and Beta Actin (Proteintech, Cat No. 66009-1-Ig).

### siRNA knock-down assay

IL-2 expanded NK cells were transfected with Accell mouse Tardbp siRNA (Dharmacon, Cat No. E-040078-00-0005) or Mdm2 siRNA (Dharmacon, Cat No. E-041098-00-0005) using Accell delivery media (Dharmacon) according to the manufacturer’s instructions in the presence of IL-15 (50 ng/ml). Accell eGFP control pool was used as siRNA control. The transfection efficiency was > 90%, measured by flow cytometry. Gene knockdown efficiency was determined by qPCR and immunoblotting. Three days after transfection, cell apoptosis and proliferation were analyzed by flow cytometry as described above.

### Luciferase reporter assay

The *Ythdf2* promoter region ranging from −2000 bp to +100 bp of the TSS was amplified from murine NK cells and cloned into a pGL4-basic luciferase reporter vector (Promega) to generate a pGL4-*Ythdf2* reporter plasmid. HEK293T cells purchased from ATCC were co-transfected with the pGL4-*Ythdf2* reporter plasmid as well as STAT5a or STAT5b overexpression plasmids or an empty vector, together with a pRL-TK Renilla reporter plasmid (Promega) for normalization of transfection efficiency. The cells were harvested for lysis 24 h after transfection, and luciferase activity was quantified fluorimetrically with the Dual Luciferase System (Promega). Primer sequences for cloning the *Ythdf2* promoter and STAT5a, STAT5b overexpression plasmids are listed in Supplemental Table 1.

### ChIP assays

ChIP assays were carried out using a Pierce Magnetic ChIP Kit (Cat No.26157, Thermo Scientific) according to the manufacturer’s instructions. Briefly, an equal amount of an anti-phospho-Stat5 (Tyr694) (Cat No. 9351, CST) or corresponding control normal rabbit IgG was separately used to precipitate the cross-linked DNA-protein complexes derived from 5 × 10^6^ purified mouse primary NK cells which were pre-treated with IL-15 (50 ng/ml) for 1 h. Following reversal of cross-linking, the DNA immunoprecipitated by the indicated Ab was tested by qPCR. The sequences of all primers are listed in Supplemental Table 1.

### *Ex vivo* cytotoxicity assay

*Ex vivo* cytotoxicity of NK cells was evaluated by standard ^51^Cr release assays. Mouse lymphoma cell lines RMA-S (MHC class I-deficient) and RMA (MHC class I-sufficient) cells, a gift of André Veillette (McGill University, Montréal, Canada), were used as target cells. Mice were treated with i.p. injection of Poly I:C (200 μg/mice) for 18h. Poly I:C-activated NK cells were isolated from the spleen using EasySep Mouse NK Cell Isolation Kit (STEMCELL Technologies). Purified NK cells were co-cultured with target cells at a ratio of 5:1, 2.5:1, and 1.25:1 in the presence of IL-2 (50 U/ml).

### m^6^A-seq

Purified splenic NK cells were expanded by IL-2 (1000 U/ml, National Cancer Institute, Cat No. Bulk Ro 23-6019).) *in vitro* for 7 days. Total RNA was isolated by TRIzol reagent (Thermo Fisher Scientific) from fifty million IL-2 expanded NK cells. Polyadenylated RNA was further enriched from total RNA by using Dynabeads^®^ mRNA Purification Kit (Invitrogen). mRNA samples were fragmented into ~100-nucleotide-long fragments with RNA fragmentation reagents (Invitrogen). Fragmented mRNA (5 μg mRNA) was used for m^6^A immunoprecipitation (m^6^A-IP) using an m^6^A antibody (Synaptic, 202003) following the standard protocol of the Magna MeRIP m^6^A Kit (Merck Millipore). RNA was enriched through RNA Clean & Concentration-5 (Zymo Research) for library generation with a KAPA RNA HyperPrep Kit (Roche). Sequencing was performed at the City of Hope Genomics Facility on an Illumina HiSeq2500 machine with single read 50 bp mode. Sequencing reads were mapped to the mouse genome using HISAT2 (Version: v101) (Kim et al., 2015). Mapped reads of IP and input libraries were provided for R package exomePeak (Meng et al., 2014). m^6^A peaks were visualized using IGV software (http://www.igv.org). The m^6^A binding motif was analyzed by MEME (http://meme-suite.org) and HOMER (http://homer.ucsd.edu/homer/motif). Called peaks were annotated by intersection with gene architecture using R package ChIPseeker (Yu et al., 2015). StringTie was used to perform expression level for mRNAs from input libraries by calculating FPKM (total exon fragments /mapped reads (millions) × exon length (kB)) (Pertea et al., 2015). The differentially expressed mRNAs were selected with log2 (fold change) >1 or log2 (fold change) <-1 and p value < 0.05 by R package edgeR (Robinson et al., 2010).

### YTHDF2 RNA immunoprecipitation sequencing (RIP-Seq)

Fifty million IL-2-expanded NK cells were harvested and washed twice with cold PBS, and the cell pellet was lysed with two volumes of lysis buffer [10 mM HEPES pH 7.6, 150 mM KCl, 2 mM EDTA, 0.5% NP-40, 0.5 mM DTT, 1:100 Protease Inhibitor cocktail (Thermo Fisher Scientific), and 400 U/mL SUPERase-In RNase Inhibitor (Thermo Fisher Scientific)]. The lysate was incubated on ice for 5 min and centrifuged for 15 min to clear the lysate. One-tenth volume of cell lysate was saved as input. The rest of the cell lysate was incubated with 5 μg anti-YTHDF2 (MBL, Cat No. RN123PW) that was pre-coupled with Protein A magnetic beads (Invitrogen, Cat: 10001D) at 4□ for 2h with gentle rotation. Afterward, the beads were washed five times with 1 ml ice-cold washing buffer (50 mM HEPES pH 7.6, 200 mM NaCl, 2 mM EDTA, 0.05% NP-40, 0.5 mM DTT, and 200 U/mL RNase inhibitor). Immunoprecipitated samples were subjected to Proteinase K digestion in wash buffer supplemented with 1% SDS and 2 mg/mL Proteinase K (Thermo Fisher Scientific) incubated with shaking at 1200 rpm at 55□ for 1 h. Total RNA was extracted from both input and immunoprecipitated RNA by adding 5 volumes of TRIzol reagent, followed by Direct-zol RNA Miniprep (Zymo Research). cDNA library generation was produced with a KAPA RNA HyperPrep Kit (Roche) and sequenced on Illumina HiSeq2500 platform. Peak calling results with IP and input libraries were generated by R package RIPSeeker (Li et al., 2013). HOMER was used to find motifs of the data distribution in peak regions. Called peaks were annotated by intersection with gene and transcript architecture using ChIPpeakAnno (Zhu et al., 2010).

### mRNA stability assay

Purified splenic NK cells from *Ythdf2*^ΔNK^ and *Ythdf2*^WT^ mice were cultured with IL-15 (50 ng/ml). Three days after culture, cells were treated with actinomycin D (5μg/ml, Sigma, Cat. A9415) for the indicated time. Cells without treatment were used as 0 h. Cells were collected at the indicated time, and total RNA was extracted from the cells for qPCR. The mRNA half-life (t_1/2_) was calculated using the method as previously described (Weng et al., 2018). Primer sequences are listed in Supplemental Table 1.

### Online database analysis

We used the online resource BioGPS (http://biogps.org) to analyze the tissue-specific expression of the *Ythdf2*. The RNA-seq datasets GSE106138, GSE113214, and GSE25672 were downloaded from the Gene Expression Omnibus (GEO, https://www.ncbi.nlm.nih.gov/geo/). Normalized data from RNA-seq analyses were exported, and the gene-expression z-score was visualized with the Heatmap.2 function within the gplots R library.

### Statistics

Unpaired Student’s t-tests (two-tailed) were performed using the Prism software. One-way or Two-way ANOVA was performed when three or more independent groups were compared. P values were adjusted for multiple comparisons using Holm-Sidak’s procedure. A *P* value of < 0.05 was considered significant. \**P* < 0.05, \*\**P* < 0.01 and \*\*\**P* < 0.001.

## Data availability

The RNA-seq, m^6^A-seq, and RIP-seq data were deposited in the Gene Expression Omnibus. The remaining data that support the findings of this study are available from the corresponding authors upon request.

## Online supplemental material

Fig. S1 shows the protein levels of YTHDF2 in immune cells, NK cells in response to IL-15 stimulation, MCMV infection, and tumor progression; the generation of mice with NK cell-specific deletion of *Ythdf2*. Fig. S2 shows the viral titers in the spleen and liver from mice infected with MCMV, the percentage and the absolute number of Ly49H^+^ NK and/or Ly49D^+^ NK cells in mice infected with MCMV. Fig. S3 shows proliferation, survival, maturation, and expression of activating and inhibitory receptors of NK cells from *Ythdf2*^WT^ and *Ythdf2*^ΔNK^ mice. Fig. S4 shows the expression of the components of IL-15 receptors, the PI3K-AKT pathway, and the MEK-ERK pathway in NK cells. Fig. S5 shows transcriptome-wide RNA-seq, m6A-seq, and RIP-seq assays in NK cells. Table S1 lists the primers used in this study.

## Acknowledgments

This work was supported by grants from the NIH (NS106170, AI129582, CA247550, CA163205, CA223400, CA068458, and CA210087), the Leukemia & Lymphoma Society (1364-19), The California Institute for Regenerative Medicine (DISC2COVID19-11947), and 2021 Exceptional Project Award from Breast Cancer Alliance.

## Authors’ contributions

S. Ma, J. Yu, and M. A. Caligiuri conceived and designed the project. S. Ma, J. Yan, and T. Barr designed and supervised experiments conducted in the laboratories. S. Ma, J. Yan, J. Zhang, and J. Yu performed experiments and/or data analyses. Z. Chen, S. Wang, J. C. Sun and J. Chen contributed reagents and material support. S. Ma, J. Yu, J. Chen, and M. A. Caligiuri wrote, reviewed and/or revised the paper. All authors discussed the results and commented on the manuscript.

---

**Figure S1. YTHDF2 expression in NK cells, deletion efficiency of *Ythdf2*^ΔNK^ mice and assessment of T cell responses in *Ythdf2*^ΔNK^ mice implanted with B16F10 tumor cells. (A)** Representative immunoblot of YTHDF2 in immune cell subsets of CD4^+^ T cells (CD3^+^CD4^+^), CD8^+^ T cells (CD3^+^CD8^+^), NK cells (CD3^-^NK1.^+^), B cells (CD3^-^NK1.1^-^CD19^+^), macrophage (CD3^-^NK1.1^-^CD19^-^CD11b^+^b4/80^-^), and DCs (CD3^-^NK1.1^-^CD19^-^CD11b^+^b4/80^-^CD11c^+^) isolated from spleen of *Ythdf2*^WT^ mice. (**B**) Representative immunoblot of m^6^A writers, readers, and erasers in NK cells isolated from the spleen of *Ythdf2*^WT^ mice stimulated with or without IL-15 (50 ng/ml) overnight. (**C**) Representative immunoblot (left) and quantified protein levels of YTHDF2 normalized to ACTIN (right) in splenic NK cells from WT and IL-15Tg mice. **(D)** RNA-seq analysis of the database GSE25672 showing the expression of m^6^A enzymes and readers in splenic NK cells during the MCMV infection. (**E**) Representative immunoblot (left) and quantified protein levels (right) of YTHDF2 in NK cells isolated from the spleen of mice infected with MCMV at the indicated time. **(F)** Representative plots showing the NK cells (CD3^-^K1.1^+^) in the lung tissues from B16F10 tumor-bearing mice on days 0, 12, and 24 post tumor injection. (**G)** Representative immunoblot (left) and quantified protein levels (right) of YTHDF2 in NK cells isolated from the lung of mice injected i.v. with B16F10 at the indicated time. (H) Schematic diagram showing the generation of mice with NK cell-specific deletion of *Ythdf2*. **(I-J)** The deletion of *Ythdf2* in NK cells from *Ythdf2*^ΔNK^ mice vs. *Ythdf2*^WT^ mice was verified by qPCR (**I**) and immunoblotting (**J**). **(K-N)** The percentages of infiltrating CD4^+^ T cells (**K**) and CD8^+^ T cells (**L**) as well as expression levels of IFN-γ (**M-N**) in the lung-infiltrated NK cells from *Ythdf2*^WT^ and *Ythdf2*^ΔNK^ mice were analyzed on day 14 after B16F10 injection (n = 5 mice per group). Each symbol represents an individual mouse. Data are shown as mean ± SD and were analyzed by unpaired two-tailed *t*-test (**C, I, K-N**), one-way ANOVA (**E, G**). Data are representative of at least two independent experiments.

**Figure S2. *Ythdf2*-deficient NK cells have impaired anti-viral functions.** (**A-B**) Viral titers in the spleen (**A**) and liver (**B**) on day 7 post-infection were analyzed by qPCR. Mice injected with PBS were used as control (n = 5 mice for MCMV groups; n = 3 for PBS groups). **(C-D)** The percentage (**C**) and the absolute number (**D**) of NK cells from *Ythdf2*^WT^ and *Ythdf2*^ΔNK^ mice in the blood on days 0, 4, and 7 post MCMV infection (n = 3 at days 0 and 4; n = 5 at day 7). **(E)** Quantification of Annexin V^+^ NK cells in spleen and blood from *Ythdf2*^WT^ and *Ythdf2*^ΔNK^ mice on day 7 post MCMV infection (n = 3 mice per group). **(F-G)** The percentage (**F**) and the absolute number (**G**) of Ly49H^+^ NK cells in blood from *Ythdf2*^WT^ and *Ythdf2*^ΔNK^ mice on days 0, 4, and 7 post MCMV infection (n = 3 at days 0 and 4; n = 5 at day 7). **(H-K)** The percentage (**H, J**) and the absolute number (**I, K**) of Ly49D^+^ NK cells in the spleen (**H, I**) and blood (**J, K**) from *Ythdf2*^WT^ and *Ythdf2*^ΔNK^ mice on days 0, 4, and 7 post MCMV infection (n = 3 at days 0 and 4; n = 5 at day 7). (**L-Q**) The percentage (**L-N**) and the absolute number (**O-Q**) of Ly49D^+^Ly49H^-^ NK cells, Ly49D^+^Ly49H^+^ NK cells, and Ly49D^-^Ly49H^+^ NK cells in the spleen from *Ythdf2*^WT^ and *Ythdf2*^ΔNK^ mice on days 0, 4, and 7 post MCMV infection (n = 3 at days 0 and 4; n = 5 at day 7). (**R-W**) The percentage (**R-T**) and the absolute number (**U-W**) of Ly49D^+^Ly49H^-^ NK cells, Ly49D^+^Ly49H^+^ NK cells, and Ly49D^-^Ly49H^+^ NK cells in the blood from *Ythdf2*^WT^ and *Ythdf2*^ΔNK^ mice on days 0, 4, and 7 post MCMV infection (n = 3 at days 0 and 4; n = 5 at day 7). **(X)** Quantification of granzyme B expression by NK cells in the spleen and blood from *Ythdf2*^WT^ and *Ythdf2*^ΔNK^ mice on day 7 post MCMV infection (n = 5 mice per group). **(Y)** Quantification of IFN-γ expression by NK cells in spleen from *Ythdf2*^WT^ and *Ythdf2*^ΔNK^ mice on days 0, 4, and 7 post MCMV infection (n = 3 at days 0, 4, 7). Data are shown as mean ± SD and were analyzed by one-way ANOVA with Sidak post-test (**A, B)**, or two-way ANOVA with Sidak post-test (**C-Y**). \**P* < 0.05, \*\**P* < 0.01, \*\*\**P* < 0.001. Data are representative of at least two independent experiments.

**Figure S3. YTHDF2 controls NK cell homeostasis, terminal maturation and effector function at a steady state. (A)** Representative plots (left) and quantification (right) of CLP (Lin^-^CD27^+^2B4^+^CD127^+^Flt3^+^CD122^-^), preNKP (Lin^-^CD27^+^2B4^+^CD127^+^lt3^-^CD122^-^), and rNKP (Lin^-^CD27^+^2B4CD127^+^lt3^-^CD122^-^) in the BM between *Ythdf2*^WT^ and *Ythdf2*^ΔNK^ mice (n = 4 per group). Cells were gated on Lin^-^CD27^+^2B4^+^CD127^+^. **(B)** The percentage of Ki67^+^ NK cells in the spleen, liver, lung, and BM from *Ythdf2*^WT^ and *Ythdf2*^ΔNK^ mice (n = 6 per group). **(C)** The percentage of Annexin V^+^ NK cells in the spleen, liver, lung, and BM from *Ythdf2*^WT^ and *Ythdf2*^ΔNK^ mice (n = 6 per group). **(D)** Scheme for evaluating NK cell homeostatic proliferation *in vivo* (left). Relative percentage (right) of transferred WT and *Ythdf2*^ΔNK^ mice in the spleen of *Rag2*^−/−^*Il2rg*^−/−^ mice at indicated time points after adoptive transfer are shown (n = 4 mice per group). **(E)** Representative histograms (left) and quantification (right) of CTV dilution of transferred WT and *Ythdf2*^ΔNK^ mice in the spleen of *Rag2*^−/−^*Il2rg*^−/−^ mice five days after adoptive transfer (n = 4 mice per group). **(F)** Representative plots (left) and quantification (right) of Annexin V^+^ NK cells of transferred WT and *Ythdf2*^ΔNK^ mice in the spleen of *Rag2*^−/−^*Il2rg*^−/−^ mice five days after adoptive transfer (n = 4 mice per group). **(G)** qPCR showing the expression *Ythdf2* in splenic NK cells with different stages. **(H)** The percentage of immature, intermediate mature, and terminal mature stages of NK cells in the blood, liver, lung, and BM from *Ythdf2*^WT^ and *Ythdf2*^ΔNK^ mice (n = 6 mice per group). **(I)** Quantification of expression levels of Tbet (left) or Eomes (right) from splenic NK cells from *Ythdf2*^WT^ and *Ythdf2*^ΔNK^ mice (n = 4 mice per group). **(J)** Representative histograms of Eomes expression in immature, intermediate mature, and terminal mature stages of splenic NK cells from *Ythdf2*^WT^ and *Ythdf2*^ΔNK^ mice. **(K-L)** qPCR showing the expression *Tbet* (**K**) or *Eomes* (**L**) in immature, intermediate mature, and terminal mature stages of splenic NK cells from *Ythdf2*^WT^ and *Ythdf2*^ΔNK^ mice. **(M)** Quantification of the indicated molecules in splenic NK cells from *Ythdf2*^WT^ and *Ythdf2*^ΔNK^ mice (n = 4 mice per group). **(N)** Cytotoxicity of NK cells against RMA cells was evaluated by standard ^51^Cr release assays (n = 3 mice per group). Data are shown as mean ± SD and were analyzed by unpaired two-tailed *t*-test (**B, C, E, F, I, M**), one-way ANOVA with Sidak post-test (**G**), or two-way ANOVA with Sidak post-test (**A, D, H, K, L, N**). \**P* < 0.05, \*\**P* < 0.01, \*\*\**P* < 0.001, \*\*\*\**P* < 0.0001. Data are representative of at least two independent experiments.

**Figure S4. *Ythdf2* deficiency has no effect on the expression of the components of IL-15 receptors, the PI3K-AKT pathway, and the MEK-ERK pathway in NK cells. (A)** qPCR showing the expression of *Ythdf2* in NK cells at indicated time points following stimulation of IL-15. **(B)** qPCR showing the expression of *Ythdf2* in NK cells in the presence of STAT5 inhibitor and IL-15 stimulation. (**C-D**) Representative histograms (left) and quantification (right) of intracellular staining showing the YTHDF2 protein levels in splenic NK cells from *Stat5*^WT^ and *Stat5*^ΔNK^ mice under the stimulation of IL-15. (**E-I**) Mice were treated with IL-15 (2 μg per day) for 5 days (n = 3 per group). The representative plots (**E**), the percentage (**F**), and the absolute number (**G**) of NK cells in the spleen are shown. (**H-J**) Representative plots of Ki67^+^ NK cells (**H**), Brdu^+^ NK cells (**I**), and Annexin V^+^ NK cells (**J**) from IL-15 treated mice are shown. **(K)** Representative histograms of CD122 and CD132 levels in splenic NK cells from *Ythdf2*^WT^ and *Ythdf2*^ΔNK^ mice. **(L-M)** Representative histograms (**L**) and quantification (**M**) of indicated molecules in splenic NK cells from *Ythdf2*^WT^ and *Ythdf2*^ΔNK^ mice after stimulation of IL-15. Data are shown as mean ± SD and were analyzed by unpaired two-tailed *t*-test (**M**), oneway ANOVA with Sidak post-test (**A, D, F, G**). \**P* < 0.01, \*\**P* < 0.01, \*\*\**P* < 0.001. Data are representative of at least two independent experiments.

**Figure S5. Transcriptome-wide RNA-seq, m^6^A-seq, and RIP-seq assays in murine NK cells. (A)** Volcano plots of differentially expressed genes from RNA-seq. (**B**) Gene Set Enrichment Analysis showing enrichment of E2F targets, G2/M checkpoint, and mitotic spindle hallmark gene sets in *Ythdf2*^ΔNK^ NK cells compared to *Ythdf2*^WT^ NK cells. **(C)** The proportion of m^6^A peak distribution in NK cells from *Ythdf2*^WT^ and *Ythdf2*^ΔNK^ mice. **(D)** GO analysis of transcripts with m^6^A peaks. **(E)** Density distribution of the YTHDF2-binding sites across mRNA transcriptome from RIP-seq data. **(F)** The proportion of YTHDF2-binding sites distribution from RIP-seq data. **(G)** Top 10 GO clusters from GO analysis of YTHDF2 target genes from RIP-seq data. (**H**) Immunoblotting showing the protein levels of MDM2, TDP-43, and YTHDF2 in IL-2 expanded splenic NK cells from *Ythdf2*^WT^ and *Ythdf2*^ΔNK^ mice. (**I-J**) qPCR and immunoblotting showing the expression of *Mdm2* (**I**) and *Tardbp* (**J**) in NK cells transfected with gene-specific or control siRNA. (**K-L**) IL-2 expanded NK cells were transfected with *Mdm2* specific siRNA cells under the stimulation of IL-15. Three days later, cell proliferation and apoptosis were analyzed by Ki67 staining (**K**) and Annexin V straining (**L**), respectively (n = 3 per group). Data are shown as mean ± SD and were analyzed by unpaired two-tailed *t*-test (**I, J**) or one-way ANOVA with Sidak post-test (**K, L**). \*\*\**P* < 0.001. Data are representative of at least two independent experiments.

## Reference

Arase, H., E.S. Mocarski, A.E. Campbell, A.B. Hill, and L.L. Lanier. 2002. Direct recognition of cytomegalovirus by activating and inhibitory NK cell receptors. Science 296:1323–1326.

Ayala, Y.M., T. Misteli, and F.E. Baralle. 2008. TDP-43 regulates retinoblastoma protein phosphorylation through the repression of cyclin-dependent kinase 6 expression. Proceedings of the National Academy of Sciences of the United States of America 105:3785–3789.

Barbieri, I., K. Tzelepis, L. Pandolfini, J. Shi, G. Millan-Zambrano, S.C. Robson, D. Aspris, V. Migliori, A.J. Bannister, N. Han, E. De Braekeleer, H. Ponstingl, A. Hendrick, C.R. Vakoc, G.S. Vassiliou, and T. Kouzarides. 2017. Promoter-bound METTL3 maintains myeloid leukaemia by m(6)A-dependent translation control. Nature 552:126–131.

Becknell, B., and M.A. Caligiuri. 2005. Interleukin-2, interleukin-15, and their roles in human natural killer cells. Adv Immunol 86:209–239.

Bertoli, C., J.M. Skotheim, and R.A. de Bruin. 2013. Control of cell cycle transcription during G1 and S phases. Nat Rev Mol Cell Biol 14:518–528.

Bezman, N.A., C.C. Kim, J.C. Sun, G. Min-Oo, D.W. Hendricks, Y. Kamimura, J.A. Best, A.W. Goldrath, L.L. Lanier, and C. Immunological Genome Project. 2012. Molecular definition of the identity and activation of natural killer cells. Nat Immunol 13:1000–1009.

Carson, W.E., T.A. Fehniger, S. Haldar, K. Eckhert, M.J. Lindemann, C.F. Lai, C.M. Croce, H. Baumann, and M.A. Caligiuri. 1997. A potential role for interleukin-15 in the regulation of human natural killer cell survival. J Clin Invest 99:937–943.

Carson, W.E., J.G. Giri, M.J. Lindemann, M.L. Linett, M. Ahdieh, R. Paxton, D. Anderson, J. Eisenmann, K. Grabstein, and M.A. Caligiuri. 1994. Interleukin (IL) 15 is a novel cytokine that activates human natural killer cells via components of the IL-2 receptor. J Exp Med 180:1395–1403.

Chan, C.J., M.J. Smyth, and L. Martinet. 2014. Molecular mechanisms of natural killer cell activation in response to cellular stress. Cell Death Differ 21:5–14.

Chen, M., L. Wei, C.T. Law, F.H. Tsang, J. Shen, C.L. Cheng, L.H. Tsang, D.W. Ho, D.K. Chiu, J.M. Lee, C.C. Wong, I.O. Ng, and C.M. Wong. 2018. RNA N6-methyladenosine methyltransferase-like 3 promotes liver cancer progression through YTHDF2-dependent posttranscriptional silencing of SOCS2. Hepatology 67:2254–2270.

Chen, X., J. Han, J. Chu, L. Zhang, J. Zhang, C. Chen, L. Chen, Y. Wang, H. Wang, L. Yi, J.B. Elder, Q.E. Wang, X. He, B. Kaur, E.A. Chiocca, and J. Yu. 2016. A combinational therapy of EGFR-CAR NK cells and oncolytic herpes simplex virus 1 for breast cancer brain metastases. Oncotarget

Chiossone, L., J. Chaix, N. Fuseri, C. Roth, E. Vivier, and T. Walzer. 2009. Maturation of mouse NK cells is a 4-stage developmental program. Blood 113:5488–5496.

Chu, J., Y. Deng, D.M. Benson, S. He, T. Hughes, J. Zhang, Y. Peng, H. Mao, L. Yi, K. Ghoshal, X. He, S.M. Devine, X. Zhang, M.A. Caligiuri, C.C. Hofmeister, and J. Yu. 2014. CS1-specific chimeric antigen receptor (CAR)-engineered natural killer cells enhance in vitro and in vivo antitumor activity against human multiple myeloma. Leukemia 28:917–927.

Cong, J., X. Wang, X. Zheng, D. Wang, B. Fu, R. Sun, Z. Tian, and H. Wei. 2018. Dysfunction of Natural Killer Cells by FBP1-Induced Inhibition of Glycolysis during Lung Cancer Progression. Cell Metab 28:243–255 e245.

Corbel, C., O. Pourquie, F. Cormier, P. Vaigot, and N.M. Le Douarin. 1996. BEN/SC1/DM-GRASP, a homophilic adhesion molecule, is required for in vitro myeloid colony formation by avian hemopoietic progenitors. Proc Natl Acad Sci U S A 93:2844–2849.

Cui, Y., G. Riedlinger, K. Miyoshi, W. Tang, C. Li, C.X. Deng, G.W. Robinson, and L. Hennighausen. 2004. Inactivation of Stat5 in mouse mammary epithelium during pregnancy reveals distinct functions in cell proliferation, survival, and differentiation. Mol Cell Biol 24:8037–8047.

Daussy, C., F. Faure, K. Mayol, S. Viel, G. Gasteiger, E. Charrier, J. Bienvenu, T. Henry, E. Debien, U.A. Hasan, J. Marvel, K. Yoh, S. Takahashi, I. Prinz, S. de Bernard, L. Buffat, and T. Walzer. 2014. T-bet and Eomes instruct the development of two distinct natural killer cell lineages in the liver and in the bone marrow. J Exp Med 211:563–577.

Deng, Y., Y. Kerdiles, J. Chu, S. Yuan, Y. Wang, X. Chen, H. Mao, L. Zhang, J. Zhang, T. Hughes, Q. Zhang, F. Wang, X. Zou, C.G. Liu, A.G. Freud, X. Li, M.A. Caligiuri, E. Vivier, and J. Yu. 2015. Transcription factor foxo1 is a negative regulator of natural killer cell maturation and function. Immunity 42:457–470.

Dixit, D., B.C. Prager, R.C. Gimple, H.X. Poh, Y. Wang, Q. Wu, Z. Qiu, R.L. Kidwell, L.J.Y. Kim, Q. Xie, K. Vitting-Seerup, S. Bhargava, Z. Dong, L. Jiang, Z. Zhu, P. Hamerlik, S.R. Jaffrey, J.C. Zhao, X. Wang, and J.N. Rich. 2020. The RNA m6A reader YTHDF2 maintains oncogene expression and is a targetable dependency in glioblastoma stem cells. Cancer Discov

Du, H., Y. Zhao, J. He, Y. Zhang, H. Xi, M. Liu, J. Ma, and L. Wu. 2016. YTHDF2 destabilizes m(6)A-containing RNA through direct recruitment of the CCR4-NOT deadenylase complex. Nat Commun 7:12626.

Fathman, J.W., D. Bhattacharya, M.A. Inlay, J. Seita, H. Karsunky, and I.L. Weissman. 2011. Identification of the earliest natural killer cell-committed progenitor in murine bone marrow. Blood 118:5439–5447.

Fauriat, C., E.O. Long, H.G. Ljunggren, and Y.T. Bryceson. 2010. Regulation of human NK-cell cytokine and chemokine production by target cell recognition. Blood 115:2167–2176.

Fehniger, T.A., K. Suzuki, A. Ponnappan, J.B. VanDeusen, M.A. Cooper, S.M. Florea, A.G. Freud, M.L. Robinson, J. Durbin, and M.A. Caligiuri. 2001. Fatal leukemia in interleukin 15 transgenic mice follows early expansions in natural killer and memory phenotype CD8_+_ T cells. J Exp Med 193:219–231.

Fei, Q., Z. Zou, I.A. Roundtree, H.L. Sun, and C. He. 2020. YTHDF2 promotes mitotic entry and is regulated by cell cycle mediators. PLoS Biol 18:e3000664.

French, A.R., H. Sjolin, S. Kim, R. Koka, L. Yang, D.A. Young, C. Cerboni, E. Tomasello, A. Ma, E. Vivier, K. Karre, and W.M. Yokoyama. 2006. DAP12 signaling directly augments proproliferative cytokine stimulation of NK cells during viral infections. J Immunol 177:4981–4990.

Frum, R., M. Ramamoorthy, L. Mohanraj, S. Deb, and S.P. Deb. 2009. MDM2 controls the timely expression of cyclin A to regulate the cell cycle. Mol Cancer Res 7: 1253–1267.

Geiger, T.L., and J.C. Sun. 2016. Development and maturation of natural killer cells. Curr Opin Immunol 39:82–89.

Giono, L.E., and J.J. Manfredi. 2007. Mdm2 is required for inhibition of Cdk2 activity by p21, thereby contributing to p53-dependent cell cycle arrest. Mol Cell Biol 27:4166–4178.

Giono, L.E., L. Resnick-Silverman, L.A. Carvajal, S. St Clair, and J.J. Manfredi. 2017. Mdm2 promotes Cdc25C protein degradation and delays cell cycle progression through the G2/M phase. Oncogene 36:6762–6773.

Gorbsky, G.J. 2015. The spindle checkpoint and chromosome segregation in meiosis. FEBS J 282:2471–2487.

Gordon, S.M., J. Chaix, L.J. Rupp, J. Wu, S. Madera, J.C. Sun, T. Lindsten, and S.L. Reiner. 2012. The transcription factors T-bet and Eomes control key checkpoints of natural killer cell maturation. Immunity 36:55–67.

Grossman, S.A., S. Ellsworth, J. Campian, A.T. Wild, J.M. Herman, D. Laheru, M. Brock, A. Balmanoukian, and X. Ye. 2015. Survival in Patients With Severe Lymphopenia Following Treatment With Radiation and Chemotherapy for Newly Diagnosed Solid Tumors. J Natl Compr Canc Netw 13:1225–1231.

Han, D., J. Liu, C. Chen, L. Dong, Y. Liu, R. Chang, X. Huang, Y. Liu, J. Wang, U. Dougherty, M.B. Bissonnette, B. Shen, R.R. Weichselbaum, M.M. Xu, and C. He. 2019. Anti-tumour immunity controlled through mRNA m(6)A methylation and YTHDF1 in dendritic cells. Nature 566:270–274.

Han, J., J. Chu, W. Keung Chan, J. Zhang, Y. Wang, J.B. Cohen, A. Victor, W.H. Meisen, S.H. Kim, P. Grandi, Q.E. Wang, X. He, I. Nakano, E.A. Chiocca, J.C. Glorioso, 3rd, B. Kaur, M.A. Caligiuri, and J. Yu. 2015. CAR-Engineered NK Cells Targeting Wild-Type EGFR and EGFRvIII Enhance Killing of Glioblastoma and Patient-Derived Glioblastoma Stem Cells. Sci Rep 5:11483.

Hou, J., H. Zhang, J. Liu, Z. Zhao, J. Wang, Z. Lu, B. Hu, J. Zhou, Z. Zhao, M. Feng, H. Zhang, B. Shen, X. Huang, B. Sun, M.J. Smyth, C. He, and Q. Xia. 2019. YTHDF2 reduction fuels inflammation and vascular abnormalization in hepatocellular carcinoma. Mol Cancer 18:163.

Hu, Z., Y. Han, Y. Liu, Z. Zhao, F. Ma, A. Cui, F. Zhang, Z. Liu, Y. Xue, J. Bai, H. Wu, H. Bian, Y.E. Chin, Y. Yu, Z. Meng, H. Wang, Y. Liu, J. Fan, X. Gao, Y. Chen, and Y. Li. 2020. CREBZF as a Key Regulator of STAT3 Pathway in the Control of Liver Regeneration in Mice. Hepatology 71:1421–1436.

Huang, H., H. Weng, W. Sun, X. Qin, H. Shi, H. Wu, B.S. Zhao, A. Mesquita, C. Liu, C.L. Yuan, Y.C. Hu, S. Huttelmaier, J.R. Skibbe, R. Su, X. Deng, L. Dong, M. Sun, C. Li, S. Nachtergaele, Y. Wang, C. Hu, K. Ferchen, K.D. Greis, X. Jiang, M. Wei, L. Qu, J.L. Guan, C. He, J. Yang, and J. Chen. 2018. Recognition of RNA N(6)-methyladenosine by IGF2BP proteins enhances mRNA stability and translation. Nat Cell Biol 20:285–295.

Kim, D., B. Langmead, and S.L. Salzberg. 2015. HISAT: a fast spliced aligner with low memory requirements. Nat Methods 12:357–360.

Larisch, S., Y. Yi, R. Lotan, H. Kerner, S. Eimerl, W. Tony Parks, Y. Gottfried, S. Birkey Reffey, M.P. de Caestecker, D. Danielpour, N. Book-Melamed, R. Timberg, C.S. Duckett, R.J. Lechleider, H. Steller, J. Orly, S.J. Kim, and A.B. Roberts. 2000. A novel mitochondrial septin-like protein, ARTS, mediates apoptosis dependent on its P-loop motif. Nat Cell Biol 2:915–921.

Lee, S.H., K.S. Kim, N. Fodil-Cornu, S.M. Vidal, and C.A. Biron. 2009. Activating receptors promote NK cell expansion for maintenance, IL-10 production, and CD8 T cell regulation during viral infection. J Exp Med 206:2235–2251.

Leong, J.W., S.E. Schneider, R.P. Sullivan, B.A. Parikh, B.A. Anthony, A. Singh, B.A. Jewell, T. Schappe, J.A. Wagner, D.C. Link, W.M. Yokoyama, and T.A. Fehniger. 2015. PTEN regulates natural killer cell trafficking in vivo. Proceedings of the National Academy of Sciences of the United States of America 112:E700–709.

Li, H.B., J. Tong, S. Zhu, P.J. Batista, E.E. Duffy, J. Zhao, W. Bailis, G. Cao, L. Kroehling, Y. Chen, G. Wang, J.P. Broughton, Y.G. Chen, Y. Kluger, M.D. Simon, H.Y. Chang, Z. Yin, and R.A. Flavell. 2017. m(6)A mRNA methylation controls T cell homeostasis by targeting the IL-7/STAT5/SOCS pathways. Nature 548:338–342.

Li, J., H. Xie, Y. Ying, H. Chen, H. Yan, L. He, M. Xu, X. Xu, Z. Liang, B. Liu, X. Wang, X. Zheng, and L. Xie. 2020. YTHDF2 mediates the mRNA degradation of the tumor suppressors to induce AKT phosphorylation in N6-methyladenosine-dependent way in prostate cancer. Mol Cancer 19:152.

Li, Y., D.Y. Zhao, J.F. Greenblatt, and Z. Zhang. 2013. RIPSeeker: a statistical package for identifying protein-associated transcripts from RIP-seq experiments. Nucleic Acids Res 41:e94.

Li, Z., P. Qian, W. Shao, H. Shi, X.C. He, M. Gogol, Z. Yu, Y. Wang, M. Qi, Y. Zhu, J.M. Perry, K. Zhang, F. Tao, K. Zhou, D. Hu, Y. Han, C. Zhao, R. Alexander, H. Xu, S. Chen, A. Peak, K. Hall, M. Peterson, A. Perera, J.S. Haug, T. Parmely, H. Li, B. Shen, J. Zeitlinger, C. He, and L. Li. 2018. Suppression of m(6)A reader Ythdf2 promotes hematopoietic stem cell expansion. Cell Res 28:904–917.

Liu, E., D. Marin, P. Banerjee, H.A. Macapinlac, P. Thompson, R. Basar, L. Nassif Kerbauy, B. Overman, P. Thall, M. Kaplan, V. Nandivada, I. Kaur, A. Nunez Cortes, K. Cao, M. Daher, C. Hosing, E.N. Cohen, P. Kebriaei, R. Mehta, S. Neelapu, Y. Nieto, M. Wang, W. Wierda, M. Keating, R. Champlin, E.J. Shpall, and K. Rezvani. 2020. Use of CAR-Transduced Natural Killer Cells in CD19-Positive Lymphoid Tumors. N Engl J Med 382:545–553.

Liu, Y., Y. You, Z. Lu, J. Yang, P. Li, L. Liu, H. Xu, Y. Niu, and X. Cao. 2019. N (6)-methyladenosine RNA modification-mediated cellular metabolism rewiring inhibits viral replication. Science 365:1171–1176.

Loh, J., D.T. Chu, A.K. O’Guin, W.M. Yokoyama, and H.W.t. Virgin. 2005. Natural killer cells utilize both perforin and gamma interferon to regulate murine cytomegalovirus infection in the spleen and liver. J Virol 79:661–667.

Lopez-Mateo, I., M.A. Villaronga, S. Llanos, and B. Belandia. 2012. The transcription factor CREBZF is a novel positive regulator of p53. Cell Cycle 11:3887–3895.

Mapperley, C., L.N. van de Lagemaat, H. Lawson, A. Tavosanis, J. Paris, J. Campos, D. Wotherspoon, J. Durko, A. Sarapuu, J. Choe, I. Ivanova, D.S. Krause, A. von Kriegsheim, C. Much, M. Morgan, R. I. Gregory, A.J. Mead, D. O’Carroll, and K.R. Kranc. 2020. The mRNA m6A reader YTHDF2 suppresses proinflammatory pathways and sustains hematopoietic stem cell function. Journal of Experimental Medicine 218:

Meng, J., Z. Lu, H. Liu, L. Zhang, S. Zhang, Y. Chen, M.K. Rao, and Y. Huang. 2014. A protocol for RNA methylation differential analysis with MeRIP-Seq data and exomePeak R/Bioconductor package. Methods 69:274–281.

Mishra, A., L. Sullivan, and M.A. Caligiuri. 2014. Molecular pathways: interleukin-15 signaling in health and in cancer. Clin Cancer Res 20:2044–2050.

Muller, J., B. Sperl, W. Reindl, A. Kiessling, and T. Berg. 2008. Discovery of chromone-based inhibitors of the transcription factor STAT5. Chembiochem 9:723–727.

Narni-Mancinelli, E., J. Chaix, A. Fenis, Y.M. Kerdiles, N. Yessaad, A. Reynders, C. Gregoire, H. Luche, S. Ugolini, E. Tomasello, T. Walzer, and E. Vivier. 2011. Fate mapping analysis of lymphoid cells expressing the NKp46 cell surface receptor. Proc Natl Acad Sci U S A 108:18324–18329.

Nguyen, K.B., T.P. Salazar-Mather, M.Y. Dalod, J.B. Van Deusen, X.Q. Wei, F.Y. Liew, M.A. Caligiuri, J.E. Durbin, and C.A. Biron. 2002. Coordinated and distinct roles for IFN-alpha beta, IL-12, and IL-15 regulation of NK cell responses to viral infection. J Immunol 169:4279–4287.

Niu, T.K., Y. Cheng, X. Ren, and J.M. Yang. 2010. Interaction of Beclin 1 with survivin regulates sensitivity of human glioma cells to TRAIL-induced apoptosis. FEBS Lett 584:3519–3524.

Orr, M.T., W.J. Murphy, and L.L. Lanier. 2010. ‘Unlicensed’ natural killer cells dominate the response to cytomegalovirus infection. Nat Immunol 11:321–327.

Orr, M.T., J.C. Sun, D.G. Hesslein, H. Arase, J.H. Phillips, T. Takai, and L.L. Lanier. 2009. Ly49H signaling through DAP10 is essential for optimal natural killer cell responses to mouse cytomegalovirus infection. J Exp Med 206:807–817.

Paris, J., M. Morgan, J. Campos, G.J. Spencer, A. Shmakova, I. Ivanova, C. Mapperley, H. Lawson, D.A. Wotherspoon, C. Sepulveda, M. Vukovic, L. Allen, A. Sarapuu, A. Tavosanis, A.V. Guitart, A. Villacreces, C. Much, J. Choe, A. Azar, L.N. van de Lagemaat, D. Vernimmen, A. Nehme, F. Mazurier, T.C.P. Somervaille, R.I. Gregory, D. O’Carroll, and K.R. Kranc. 2019. Targeting the RNA m(6)A Reader YTHDF2 Selectively Compromises Cancer Stem Cells in Acute Myeloid Leukemia. Cell Stem Cell 25:137–148 e136.

Pertea, M., G.M. Pertea, C.M. Antonescu, T.C. Chang, J.T. Mendell, and S.L. Salzberg. 2015. StringTie enables improved reconstruction of a transcriptome from RNA-seq reads. Nat Biotechnol 33:290–295.

Reiter, Z. 1993. Interferon--a major regulator of natural killer cell-mediated cytotoxicity. J Interferon Res 13:247–257.

Robinson, M.D., D.J. McCarthy, and G.K. Smyth. 2010. edgeR: a Bioconductor package for differential expression analysis of digital gene expression data. Bioinformatics 26:139–140.

Rubio, R.M., D.P. Depledge, C. Bianco, L. Thompson, and I. Mohr. 2018. RNA m(6) A modification enzymes shape innate responses to DNA by regulating interferon beta. Genes Dev 32:1472–1484.

Ruggeri, L., M. Capanni, E. Urbani, K. Perruccio, W.D. Shlomchik, A. Tosti, S. Posati, D. Rogaia, F. Frassoni, F. Aversa, M.F. Martelli, and A. Velardi. 2002. Effectiveness of donor natural killer cell alloreactivity in mismatched hematopoietic transplants. Science 295:2097–2100.

Sanna, S., S. Esposito, A. Masala, P. Sini, G. Nieddu, M. Galioto, M. Fais, C. Iaccarino, G. Cestra, and C. Crosio. 2020. HDAC1 inhibition ameliorates TDP-43-induced cell death in vitro and in vivo. Cell Death Dis 11:369.

Santana Carrero, R.M., F. Beceren-Braun, S.C. Rivas, S.M. Hegde, A. Gangadharan, D. Plote, G. Pham, S.M. Anthony, and K.S. Schluns. 2019. IL-15 is a component of the inflammatory milieu in the tumor microenvironment promoting antitumor responses. Proceedings of the National Academy of Sciences of the United States of America 116:599–608.

Shi, H., J. Wei, and C. He. 2019. Where, When, and How: Context-Dependent Functions of RNA Methylation Writers, Readers, and Erasers. Mol Cell 74:640–650.

Shulman, Z., and N. Stern-Ginossar. 2020. The RNA modification N(6)-methyladenosine as a novel regulator of the immune system. Nat Immunol 21:501–512.

Smith, H.R., J.W. Heusel, I.K. Mehta, S. Kim, B.G. Dorner, O.V. Naidenko, K. Iizuka, H. Furukawa, D.L. Beckman, J.T. Pingel, A.A. Scalzo, D.H. Fremont, and W.M. Yokoyama. 2002. Recognition of a virus-encoded ligand by a natural killer cell activation receptor. Proceedings of the National Academy of Sciences of the United States of America 99:8826–8831.

Spits, H., J.H. Bernink, and L. Lanier. 2016. NK cells and type 1 innate lymphoid cells: partners in host defense. Nat Immunol 17:758–764.

Sumaria, N., S.L. van Dommelen, C.E. Andoniou, M.J. Smyth, A.A. Scalzo, and M.A. Degli-Esposti. 2009. The roles of interferon-gamma and perforin in antiviral immunity in mice that differ in genetically determined NK-cell-mediated antiviral activity. Immunol Cell Biol 87:559–566.

Sun, J.C., J.N. Beilke, N.A. Bezman, and L.L. Lanier. 2011. Homeostatic proliferation generates long-lived natural killer cells that respond against viral infection. J Exp Med 208:357–368.

Sun, J.C., and L.L. Lanier. 2011. NK cell development, homeostasis and function: parallels with CD8(+) T cells. Nat Rev Immunol 11:645–657.

Tang, X., L. Yang, Z. Li, A.P. Nalin, H. Dai, T. Xu, J. Yin, F. You, M. Zhu, W. Shen, G. Chen, X. Zhu, D. Wu, and J. Yu. 2018. First-in-man clinical trial of CAR NK-92 cells: safety test of CD33-CAR NK-92 cells in patients with relapsed and refractory acute myeloid leukemia. Am J Cancer Res 8:1083–1089.

Tong, J., G. Cao, T. Zhang, E. Sefik, M.C. Amezcua Vesely, J.P. Broughton, S. Zhu, H. Li, B. Li, L. Chen, H.Y. Chang, B. Su, R.A. Flavell, and H.B. Li. 2018. m(6)A mRNA methylation sustains Treg suppressive functions. Cell Res 28:253–256.

Vu, L.P., B.F. Pickering, Y. Cheng, S. Zaccara, D. Nguyen, G. Minuesa, T. Chou, A. Chow, Y. Saletore, M. MacKay, J. Schulman, C. Famulare, M. Patel, V.M. Klimek, F.E. Garrett-Bakelman, A. Melnick, M. Carroll, C.E. Mason, S.R. Jaffrey, and M.G. Kharas. 2017. The N(6)-methyladenosine (m(6)A)-forming enzyme METTL3 controls myeloid differentiation of normal hematopoietic and leukemia cells. Nat Med 23:1369–1376.

Wang, H., X. Hu, M. Huang, J. Liu, Y. Gu, L. Ma, Q. Zhou, and X. Cao. 2019a. Mettl3-mediated mRNA m(6)A methylation promotes dendritic cell activation. Nat Commun 10:1898.

Wang, H., H. Zuo, J. Liu, F. Wen, Y. Gao, X. Zhu, B. Liu, F. Xiao, W. Wang, G. Huang, B. Shen, and Z. Ju. 2018a. Loss of YTHDF2-mediated m(6)A-dependent mRNA clearance facilitates hematopoietic stem cell regeneration. Cell Res 28:1035–1038.

Wang, X., Z. Lu, A. Gomez, G.C. Hon, Y. Yue, D. Han, Y. Fu, M. Parisien, Q. Dai, G. Jia, B. Ren, T. Pan, and C. He. 2014. N6-methyladenosine-dependent regulation of messenger RNA stability. Nature 505:117–120.

Wang, X., B.S. Zhao, I.A. Roundtree, Z. Lu, D. Han, H. Ma, X. Weng, K. Chen, H. Shi, and C. He. 2015. N(6)-methyladenosine Modulates Messenger RNA Translation Efficiency. Cell 161:1388–1399.

Wang, Y., J. Chu, P. Yi, W. Dong, J. Saultz, Y. Wang, H. Wang, S. Scoville, J. Zhang, L.C. Wu, Y. Deng, X. He, B. Mundy-Bosse, A.G. Freud, L.S. Wang, M.A. Caligiuri, and J. Yu. 2018b. SMAD4 promotes TGF-beta-independent NK cell homeostasis and maturation and antitumor immunity. J Clin Invest 128:5123–5136.

Wang, Y., Y. Zhang, P. Yi, W. Dong, A.P. Nalin, J. Zhang, Z. Zhu, L. Chen, D.M. Benson, B.L. Mundy-Bosse, A.G. Freud, M.A. Caligiuri, and J. Yu. 2019b. The IL-15-AKT-XBP1s signaling pathway contributes to effector functions and survival in human NK cells. Nat Immunol 20:10–17.

Weng, H., H. Huang, H. Wu, X. Qin, B.S. Zhao, L. Dong, H. Shi, J. Skibbe, C. Shen, C. Hu, Y. Sheng, Y. Wang, M. Wunderlich, B. Zhang, L.C. Dore, R. Su, X. Deng, K. Ferchen, C. Li, M. Sun, Z. Lu, X. Jiang, G. Marcucci, J.C. Mulloy, J. Yang, Z. Qian, M. Wei, C. He, and J. Chen. 2018. METTL14 Inhibits Hematopoietic Stem/Progenitor Differentiation and Promotes Leukemogenesis via mRNA m(6)A Modification. Cell Stem Cell 22:191–205 e199.

Wiedemann, G.M., S. Grassmann, C.M. Lau, M. Rapp, A.V. Villarino, C. Friedrich, G. Gasteiger, J.J. O’Shea, and J.C. Sun. 2020. Divergent Role for STAT5 in the Adaptive Responses of Natural Killer Cells. Cell Rep 33:108498.

Winkler, R., E. Gillis, L. Lasman, M. Safra, S. Geula, C. Soyris, A. Nachshon, J. Tai-Schmiedel, N. Friedman, V.T.K. Le-Trilling, M. Trilling, M. Mandelboim, J.H. Hanna, S. Schwartz, and N. Stern-Ginossar. 2019. m(6)A modification controls the innate immune response to infection by targeting type I interferons. Nat Immunol 20:173–182.

Wu, C., W. Chen, J. He, S. Jin, Y. Liu, Y. Yi, Z. Gao, J. Yang, J. Yang, J. Cui, and W. Zhao. 2020a. Interplay of m(6)A and H3K27 trimethylation restrains inflammation during bacterial infection. Sci Adv 6:eaba0647.

Wu, C., I. Macleod, and A.I. Su. 2013. BioGPS and MyGene.info: organizing online, gene-centric information. Nucleic Acids Res 41:D561–565.

Wu, S.Y., T. Fu, Y.Z. Jiang, and Z.M. Shao. 2020b. Natural killer cells in cancer biology and therapy. Mol Cancer 19:120.

Yang, W., L.M. Rozan, E.R. McDonald, 3rd, A. Navaraj, J.J. Liu, E.M. Matthew, W. Wang, D.T. Dicker, and W.S. El-Deiry. 2007. CARPs are ubiquitin ligases that promote MDM2-independent p53 and phospho-p53ser20 degradation. J Biol Chem 282:3273–3281.

Yilmaz, A., H. Cui, M.A. Caligiuri, and J. Yu. 2020. Chimeric antigen receptor-engineered natural killer cells for cancer immunotherapy. J Hematol Oncol 13:168.

Yoon, K.J., F.R. Ringeling, C. Vissers, F. Jacob, M. Pokrass, D. Jimenez-Cyrus, Y. Su, N.S. Kim, Y. Zhu, L. Zheng, S. Kim, X. Wang, L.C. Dore, P. Jin, S. Regot, X. Zhuang, S. Canzar, C. He, G.L. Ming, and H. Song. 2017. Temporal Control of Mammalian Cortical Neurogenesis by m(6)A Methylation. Cell 171:877–889 e817.

Yu, G., L.G. Wang, and Q.Y. He. 2015. ChIPseeker: an R/Bioconductor package for ChIP peak annotation, comparison and visualization. Bioinformatics 31:2382–2383.

Yu, J., A.G. Freud, and M.A. Caligiuri. 2013. Location and cellular stages of natural killer cell development. Trends Immunol 34:573–582.

Yu, J., M. Wei, B. Becknell, R. Trotta, S. Liu, Z. Boyd, M.S. Jaung, B.W. Blaser, J. Sun, D.M. Benson, Jr., H. Mao, A. Yokohama, D. Bhatt, L. Shen, R. Davuluri, M. Weinstein, G. Marcucci, and M.A. Caligiuri. 2006. Pro- and antiinflammatory cytokine signaling: reciprocal antagonism regulates interferon-gamma production by human natural killer cells. Immunity 24:575–590.

Yue, Y., J. Liu, and C. He. 2015. RNA N6-methyladenosine methylation in post-transcriptional gene expression regulation. Genes Dev 29:1343–1355.

Zhang, C., S. Huang, H. Zhuang, S. Ruan, Z. Zhou, K. Huang, F. Ji, Z. Ma, B. Hou, and X. He. 2020. YTHDF2 promotes the liver cancer stem cell phenotype and cancer metastasis by regulating OCT4 expression via m6A RNA methylation. Oncogene 39:4507–4518.

Zhao, Q., M. Meng, R. Kumar, Y. Wu, J. Huang, Y. Deng, Z. Weng, and L. Yang. 2020. Lymphopenia is associated with severe coronavirus disease 2019 (COVID-19) infections: A systemic review and meta-analysis. Int J Infect Dis 96:131–135.

Zheng, Z., L. Zhang, X.L. Cui, X. Yu, P.J. Hsu, R. Lyu, H. Tan, M. Mandal, M. Zhang, H.L. Sun, A. Sanchez Castillo, J. Peng, M.R. Clark, C. He, and H. Huang. 2020. Control of Early B Cell Development by the RNA N(6)-Methyladenosine Methylation. Cell Rep 31:107819.

Zhou, Z., H. He, K. Wang, X. Shi, Y. Wang, Y. Su, Y. Wang, D. Li, W. Liu, Y. Zhang, L. Shen, W. Han, L. Shen, J. Ding, and F. Shao. 2020. Granzyme A from cytotoxic lymphocytes cleaves GSDMB to trigger pyroptosis in target cells. Science 368:

Zhu, L.J., C. Gazin, N.D. Lawson, H. Pages, S.M. Lin, D.S. Lapointe, and M.R. Green. 2010. ChIPpeakAnno: a Bioconductor package to annotate ChIP-seq and ChIP-chip data. BMC Bioinformatics 11:237.

