## Supplementary figures and images for "The RNA m^6^A reader YTHDF2 controls NK cell anti-tumor and anti-viral immunity"

### Figure S1

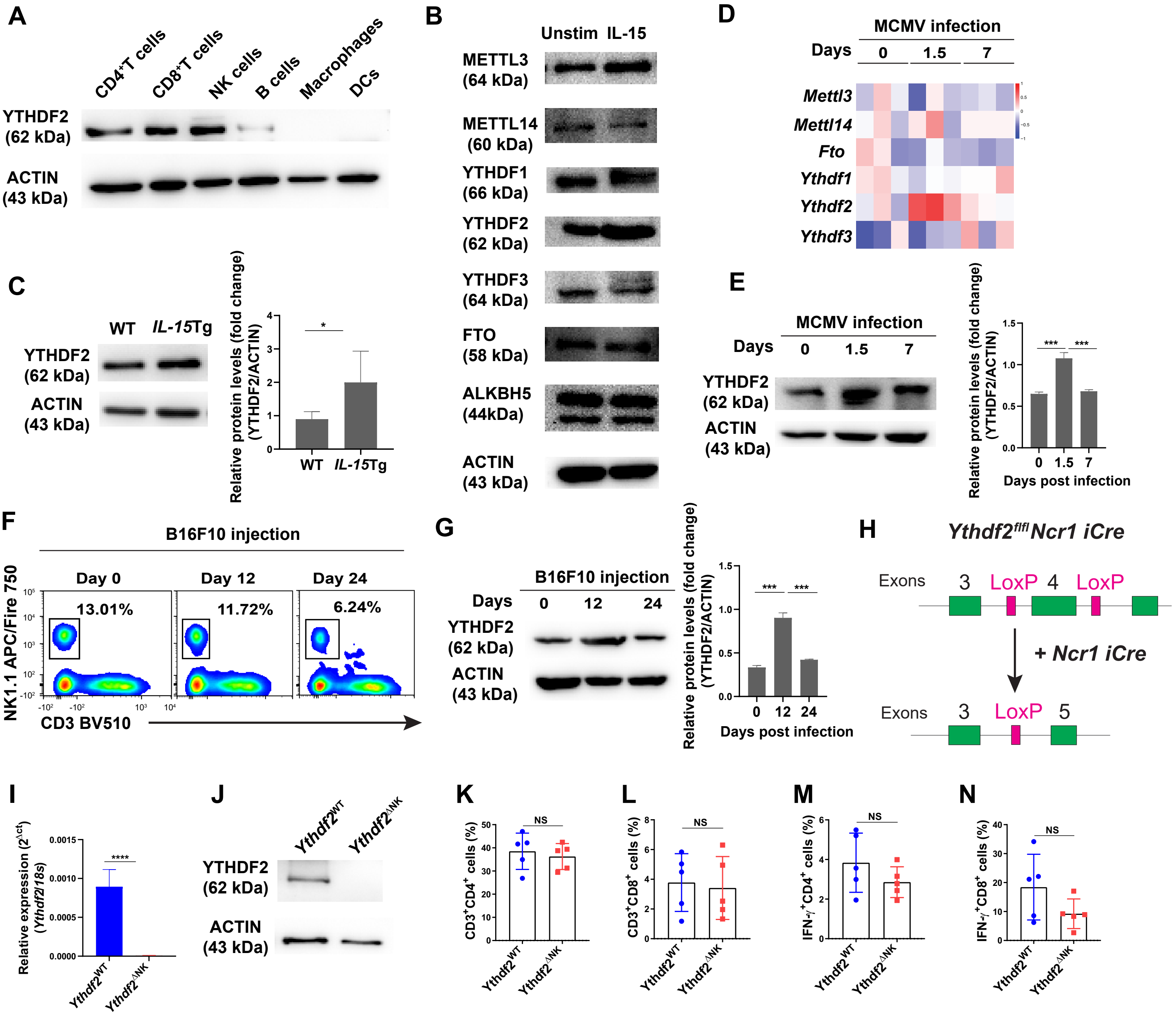

### Figure S2

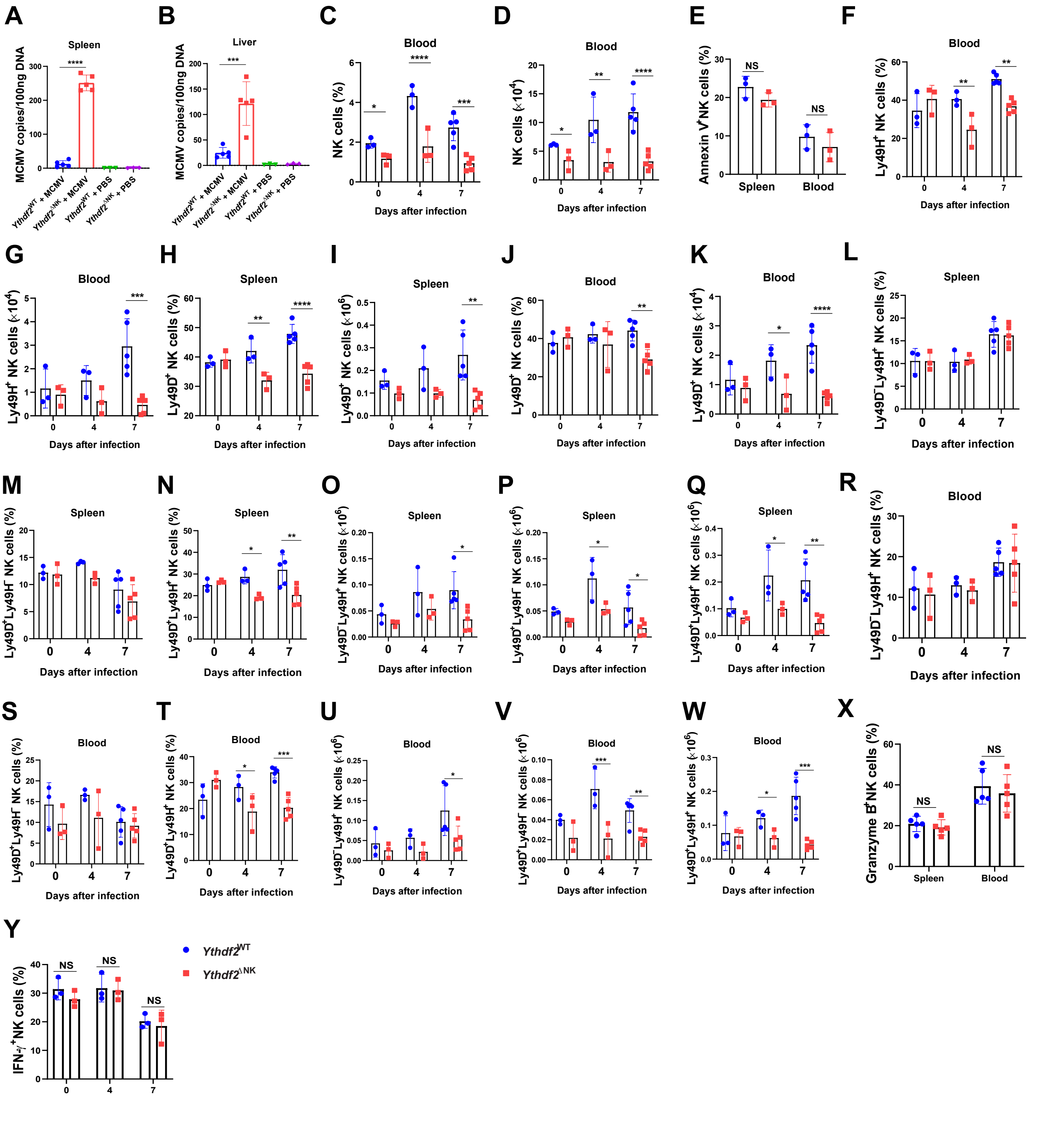

### Figure S3

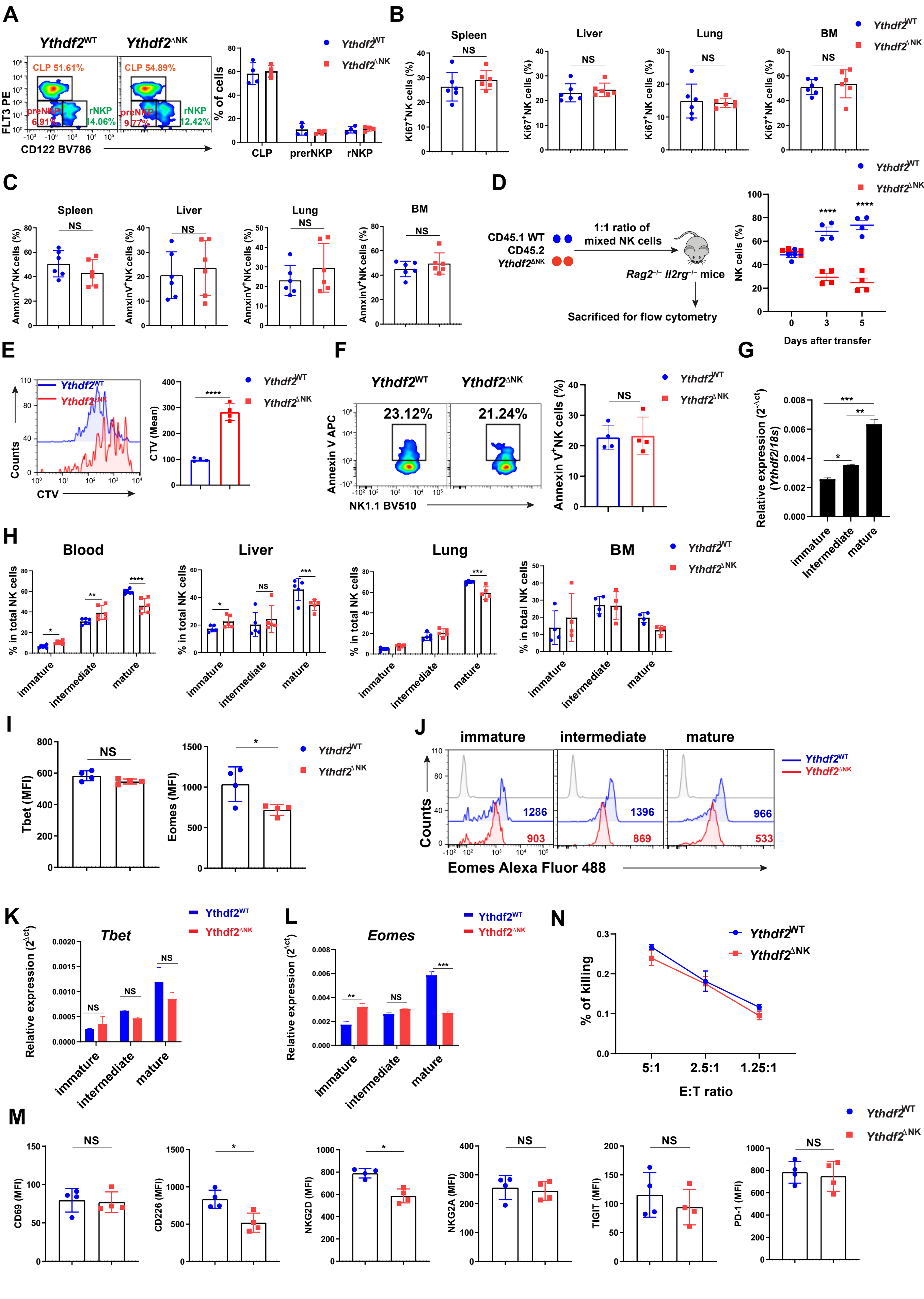

### Figure S4

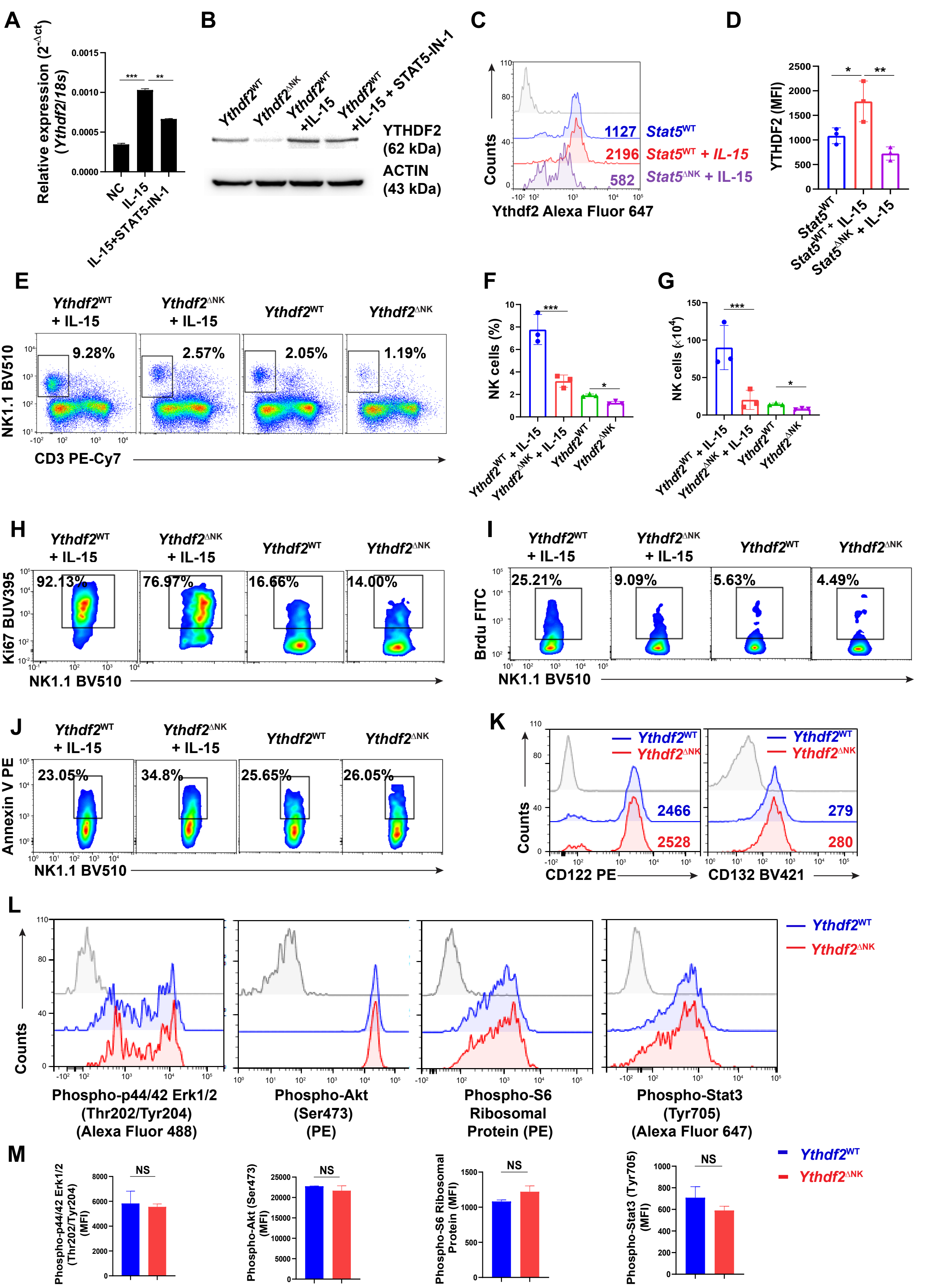

### Figure S5

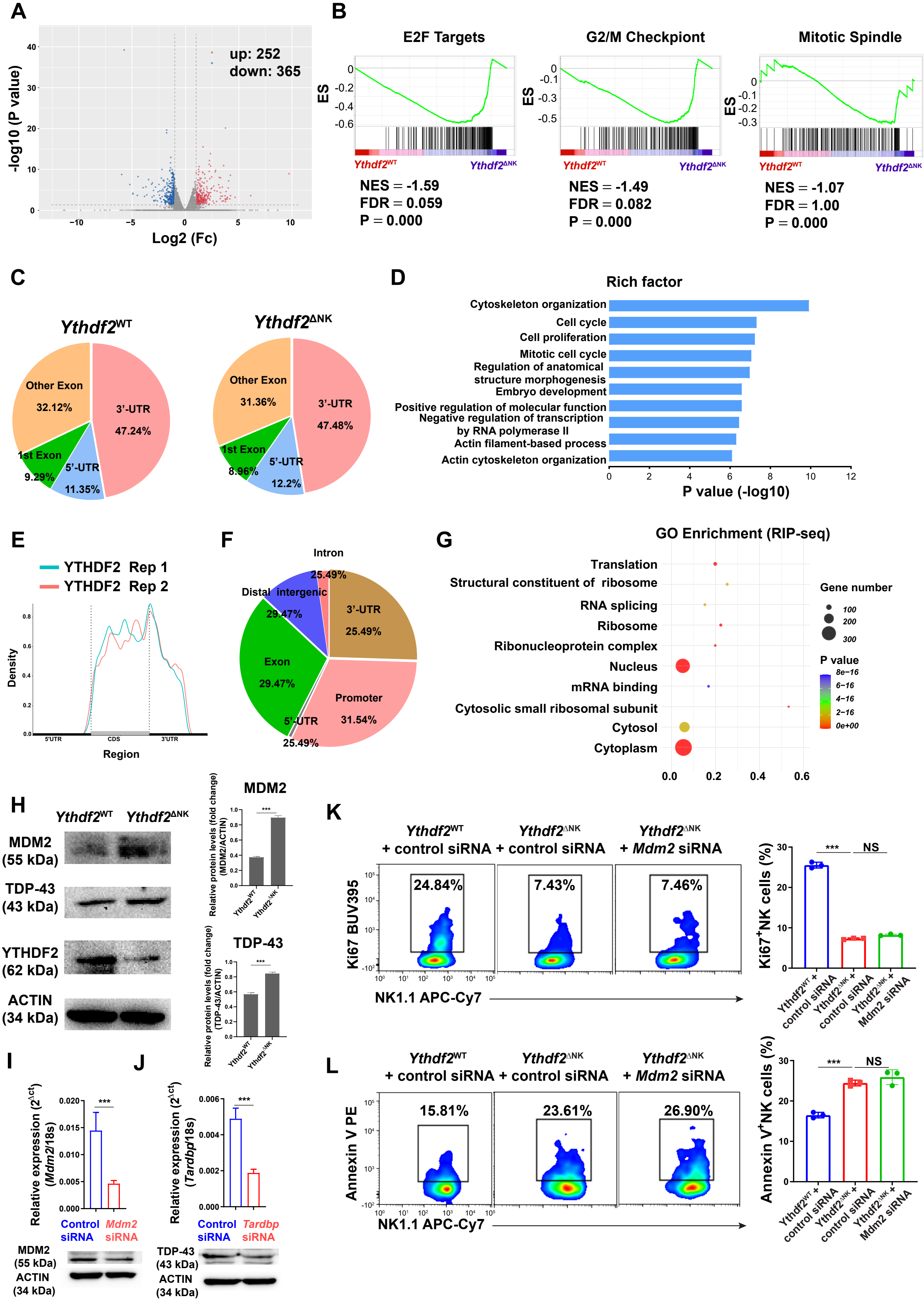
