## Supplementary material for "The RNA m^6^A reader YTHDF2 controls NK cell anti-tumor and anti-viral immunity": Table S1

**Table S1: List of oligonucleotides**

| **Name** | **Sequence** | **Note** |
| --- | --- | --- |
| mMettl3-qPCR-F | CTGGGCACTTGGATTTAAGGAA | qPCR of mouse genes |
| mMettl3-qPCR-R | TGAGAGGTGGTGTAGCAACTT |  |
| mMettl14-qPCR-F | CTGAGAGTGCGGATAGCATTG |  |
| mMettl14-qPCR-R | GAGCAGATGTATCATAGGAAGCC |  |
| mFto-qPCR-F | TCACAGCCTCGGTTTAGTTC |  |
| mFto-qPCR-R | GCAGGATCAAAGGATTTCAACG |  |
| mAlkbh5-qPCR-F | AGTTCCAGTTCAAGCCCATC |  |
| mAlkbh5-qPCR-R | GGCGTTCCTTAATGTCCTGAG |  |
| mYthdf1-qPCR-F | ACAGTTACCCCTCGATGAGTG |  |
| mYthdf1-qPCR-R | GGTAGTGAGATACGGGATGGGA |  |
| mYthdf2-qPCR-F | GAGCAGAGACCAAAAGGTCAAG |  |
| mYthdf2-qPCR-R | CTGTGGGCTCAAGTAAGGTTC |  |
| mYthdf3-qPCR-F | CATAGGGCAACAGAGGAAACAG |  |
| mYthdf3-qPCR-R | ATCTCCAGCCGTGGACCAT |  |
| mIfng-qPCR-F | ATGAACGCTACACACTGCATC |  |
| mIfng-qPCR-R | CCATCCTTTTGCCAGTTCCTC |  |
| mGzmb-qPCR-F | CCACTCTCGACCCTACATGG |  |
| mGzmb-qPCR-R | GGCCCCCAAAGTGACATTTATT |  |
| mPrf1-qPCR-F | AGCACAAGTTCGTGCCAGG |  |
| mPrf1-qPCR-R | GCGTCTCTCATTAGGGAGTTTTT |  |
| mMdm2-qPCR-F | TGTCTGTGTCTACCGAGGGTG |  |
| mMdm2-qPCR-R | TCCAACGGACTTTAACAACTTCA |  |
| mCrebzf-qPCR-F | CTGCCCGTCTTAATCGGCTC |  |
| mCrebzf-qPCR-R | CCGTAGGTAGCGACTCTCCTC |  |
| mTardbp-qPCR-F | AATCAGGGTGGGTTTGGTAACA |  |
| mTardbp-qPCR-R | GCTGGGTTAATGCTAAAAGCAC |  |
| Mdm2-m6A-F | GGGCACCTGATGCTGATCTCT | m6A-qPCR and RIP-qPCR |
| Mdm2-m6A-R | GGGTGTATGTAGTACATCCATGTG |  |
| Tardbp-m6A-F | CCCGCCTAGCGTTTATTTTTG |  |
| Tardbp-m6A-R | GAGCTGGGAGAGCAATGTCTG |  |
| Crebzf-m6A-F | GGGAGGAGTGTGTCTCCATGT |  |
| Crebzf-m6A-R | TTTTCCGCCTCTGAAATGCGT |  |
| Ythdf2 site 1-F | AGCGTTCATTTTCCAGTCCTT | CHIP-qPCR |
| Ythdf2 site 1-R | AAACACCCAGGAGCAGTACTT |  |
| Ythdf2 site 2-F | AGGGTCCAACAGACTTCAAGA |  |
| Ythdf2 site 2-R | CCTGCACACAAGACATTTACC |  |
| Ythdf2 site 3-F | ATTAAAGGCGAGCGCCACCAC |  |
| Ythdf2 site 3-R | ACATCATCTCCTGAGAACGAG |  |
| Ythdf2 site 4-F | TCAAGTCCCTGAGGAAGGAAG |  |
| Ythdf2 site 4-R | TAGGACTTGAGCCCTAACTCA |  |
| Ythdf2 NC-F | TTGGTCTGCCAATAAAAGGTT |  |
| Ythdf2 NC-R | ACTGAGGAGTGAGCTGTAGCC |  |
| Ythdf2-FRT-tF2 | ACCACATATGTGCATTGCTCACAGA | Genotyping of *Ythdf2* ^f/f^ mice |
| Ythdf2-FRT-tR2 | GCCTCCTGCCTCACTTCTAAGTGTT |  |
| MCMV-IE1-F | AGCCACCAACATTGACCACGCAC | qPCR of MCMV gene |
| MCMV-IE1-R | GCCCCAACCAGGACACACAACTC |  |
| pCDH-STAT5a-F: | CTAGTCTAGAGCCACCATGGCGGGCTGG  ATTCAGGCCCAGCA | STAT5 overexpressing plasmids |
| pCDH-STAT5a-F: | TTTTCCTTTTGCGGCCGCTCAGGACAGGG  AGCTTCTAGCGGA |  |
| pCDH-STAT5b-F | CTAGTCTAGAGCCACCATGGCTATGTGGATACAGGCTCAGCA |  |
| pCDH-STAT5b-R | TTTTCCTTTTGCGGCCGCCATGACTG  TGCGTGAGGGATCCAC |  |
| PGL4-Ythdf2-F | CGGGGTACCGTTAATAACTCATGGTCCTTC | Ythdf2 promoter |
| PGL4-Ythdf2-R | CCCAAGCTTCTCAACGCCCGTCTCTAT |  |
